# Transitions in the proteome and phospho-proteome during *Xenopus laevis* development

**DOI:** 10.1101/2021.08.05.455309

**Authors:** Elizabeth Van Itallie, Matthew Sonnett, Marian Kalocsay, Martin Wühr, Leonid Peshkin, Marc W. Kirschner

**Affiliations:** Department of Systems Biology, Harvard Medical School, Boston, MA 02115; Laboratory of Systems Pharmacology, Harvard Medical School, Boston, MA 02115; Department of Molecular Biology, Princeton University, Princeton, NJ 08544; The Lewis-Sigler Institute for Integrative Genomics, Princeton University, Princeton, NJ 08544; Eugene Bell Center, Marine Biological Laboratory, Woods Hole, MA 02543; Duke Human Vaccine Institute, Duke University School of Medicine, Durham, NC 27710; Department of Experimental Radiation Oncology, The University of Texas MD Anderson Cancer Center, Houston, TX 77030

## Abstract

Vertebrate development from an egg to a complex multi-cell organism is accompanied by multiple phases of genome-scale changes in the repertoire of proteins and their post-translational modifications. While much has been learned at the RNA level, we know less about changes at the protein level. In this paper, we present a deep analysis of changes of ∼15,000 proteins and ∼11,500 phospho-sites at 11 developmental time points in *Xenopus laevis* embryos ranging from the stage VI oocyte to juvenile tadpole. We find that the most dramatic changes to the proteome occur during the transition to functional organ systems, which occurs as the embryo becomes a tadpole. At that time, the absolute amount of non-yolk protein increases two-fold, and there is a shift in the balance of expression from proteins regulating gene expression to receptors, ligands, and proteins involved in cell-cell and cell-environment interactions. Between the early and late tadpole, the median increase for membrane and secreted proteins is substantially higher than that of nuclear proteins. To begin to appreciate changes at the post-translational level, we have measured quantitative phospho-proteomic data across the same developmental stages. In contrast to the significant protein changes that are concentrated at the end of the time series, the most significant phosphorylation changes are concentrated in the very early stages of development. A clear exception are phosphorylations of proteins involved in gene expression: these increase just after fertilization, with patterns that are highly correlated with the underlying protein changes. To facilitate the interpretation of this unique phospho-protoeme data set, we created a pipeline for identifying homologous human phosphorylations from the measured Xenopus phospho-proteome. Overall, we detected many profound temporal transitions, that suggest concerted changes in developmental strategies in the embryo, which are particularly pronounced once early patterning and specification are complete.

## INTRODUCTION

How an egg develops into an organism is a fundamental question that was addressed by morphology and comparative anatomy in the 19^th^ and early 20^th^ centuries and from a molecular genetic viewpoint in the late 20^th^ century, when many regulatory genes and pathways were identified. More recently, transcriptional profiling has revealed developmental trajectories at the level of gene expression. RNA transcription is clearly related to developmental transitions, but more and more we realize that measurement of protein abundance and posttranslational modifications will be needed to build a molecular picture of developmental regulation.

Mass spectrometry-based proteomics with isobaric tags and MS^3^ measurement can quantify many proteins accurately across different timepoints [1]. *Xenopus laevis* eggs and embryos are an ideal model for application of this technology to embryogenesis, because they are large (1.2 mm diameter), the yolk can be quantitatively removed, a single egg has 25 ug of non-yolk protein, and large numbers of staged samples can be collected easily [2–6]. Previously, we reported protein levels across development from the egg through the tailbud stage [6,7] and found temporal trajectories of most proteins to be almost flat across this period. However, those data only reported on ∼6,000 proteins at six timepoints with no replicates [7]. Additionally, the measured time period did not extend to the juvenile tadpole. Studies of recruitment of ribosomes into polysomes [8], translation of ribosomal protein mRNA [9], and yolk consumption [10] predict that the greatest amount of protein mass increase occurs between the late tailbud embryo and juvenile tadpole. Hence, we now greatly extend the time course, and also the number of proteins quantified, to report the first full proteome of a developing embryo.

To understand properly how the proteome directs development, we need to measure not just the amount of all proteins, but also their state of biological activity. The most pervasive single measurement we could do to reflect changes in protein activity or stability is to measure protein phosphorylation. Almost half the human proteome is phosphorylated [11], making it by far the most significant post-translational modification for regulating protein activity. Our lab and others previously reported phospho-proteomes during oocyte maturation through the first cleavage, which are dominated by cell cycle changes and events of meiosis [3,12]. However, these studies are not informative for the role of protein phosphorylation in embryonic development. One study included gastrula stage embryos for identification of phosphorylations [13], but no quantitative studies of the phospho-proteome after the cleavage stages have been published. However, measuring and interpreting phospho-proteomics data from a complex embryo has challenges well beyond those encountered in protein level proteomics. These include the heterogeneity of both protein and phosphorylation level differences across the embryo, the limitations in detection and identification of phosphorylated peptides [14,15], and the dearth of information on how phosphorylation of a specific residue changes protein function or localization [16]. We therefore had no expectation of what we would learn from phospho-proteomics of a developing embryo.

This paper presents the most comprehensive measurement of the proteome and phospho-proteome during embryogenesis to date. We have tried to provide this in a form of an accessible resource, which we hope will stimulate further studies of the protein level correlates of developmental processes, which have heretofore been characterized principally at the anatomical and transcriptional level.

## RESULTS

### Data Collection

We measured the proteome and phospho-proteome of *Xenopus laevis* embryogenesis across multiple developmental periods at the stages indicated in *Figure 1A*. We performed the full analysis in triplicate, collecting embryos from three independent clutches. Digested peptides were labeled with tandem mass tag reagents (TMT) and combined for enrichment of phosphorylated peptides using immobilized metal affinity chromatography (*Methods*) [14]. For two replicates, the measurements of the phospho-enriched and phospho-depleted peptides were made with Multi-Notch MS^3^ quantification (*Methods*) [1,17]. For the third replicate, the measurements of the phospho-enriched and pre-enrichment peptides were made with MS^2^ quantification.

**Figure 1.**
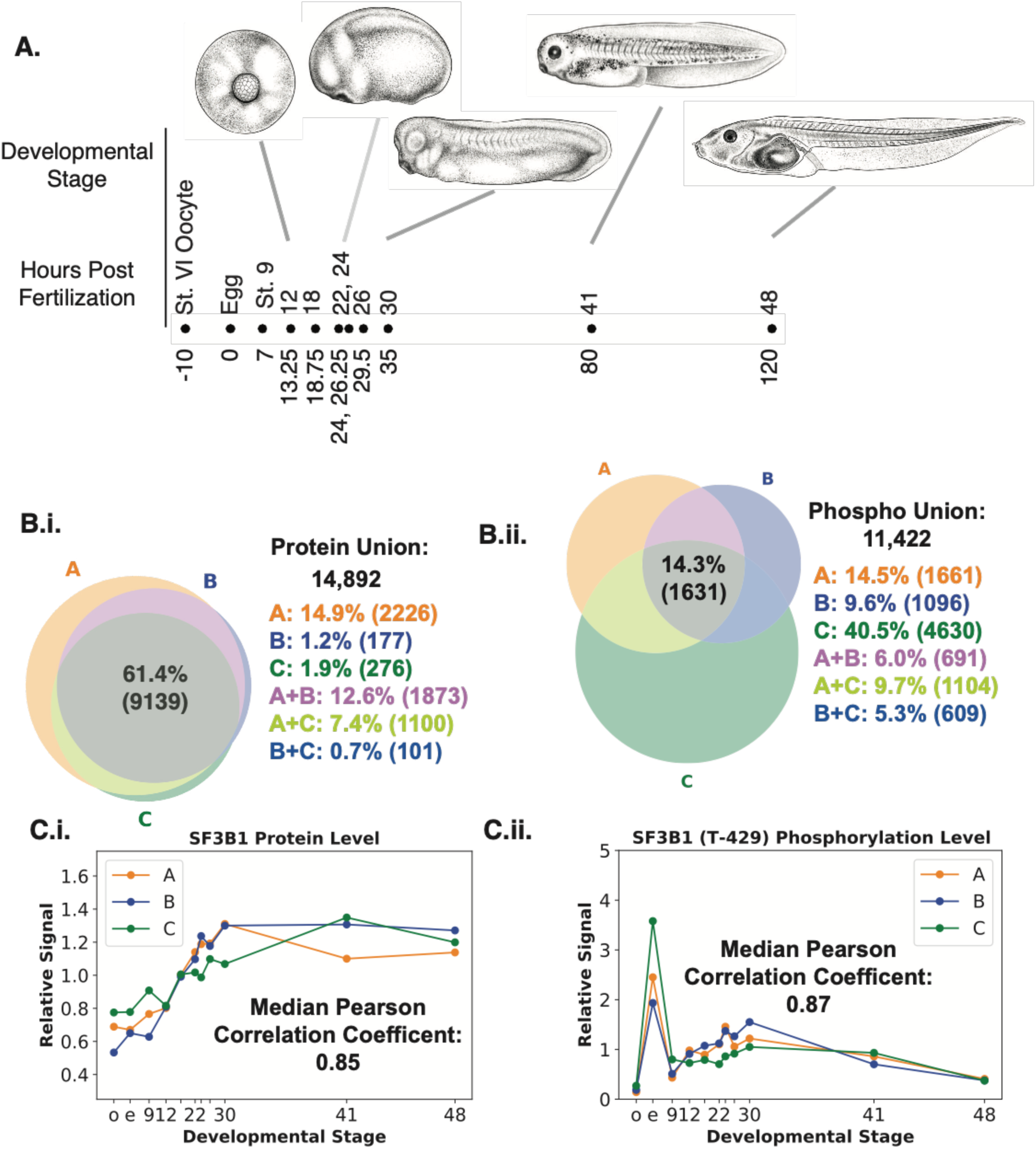
Temporal Profiling of the *Xenopus laevis* Proteome and Phospho-proteome Across Development. **A.** The eleven *Xenopus laevis* developmental stages collected for proteomic and phospho-proteomic measurement spaced according to Nieuwkoop and Faber times for temperatures between 22-24℃ and mapped to broad developmental classifications. (The Stage 41 illustration is actually Stage 40 for convenience. See Methods for Illustration information.) **B. (i)** 14,892 proteins were quantified; 61.4% measured in all three replicates. **(ii)** 11,422 phospho-forms were quantified; 14.3% were measured in all three replicates. **C.** Representative Examples: **(i)** SF3B1 protein and **(ii)** SF3B1 phospho-form T-429 are measured in all three replicates. SF3B1 replicate Pearson correlation coefficients (0.85 for protein and 0.87 for phospho-form) align with the median values for all measurements (Median Pearson of 0.90 for 14.9k proteins and 0.87 for 11.4k phospho-forms respectively; FigS2.C).

### Data Reliability

After mapping peptides to the *Xenopus laevis* reference version 9.2 and considering all measurements together when controlling for false discovery (2%), we measured a union of 14,892 proteins and 11,422 phospho-forms with 61% and 14%, respectively, measured in all replicates, and 72% and 35%, respectively, measured in at least two replicates (*Figure 1.B.i,ii*; *Methods*). Phosphorylations were measured on a union of 4,541 proteins. This number is smaller than the number of phospho-forms because 50% of phospho-proteins are measured with more than one phospho-form (*Figure S1*). Ninety-four percent of the union of phospho-proteins also have a protein measurement in at least one replicate. For replicates A, B, and C, 98%, 93%, and 86%, respectively, of phospho-form protein references also have a protein measurement for that same replicate. Thus, we measured the proteome with sufficient depth to interpret the phospho-proteome measurements.

We next assessed the reproducibility of our replicates. For each pair of replicates, we computed the Pearson Correlation Coefficient (PCC) for each timepoint in one replicate against all timepoints in the other replicate for each measured protein and phospho-form. For replicates A and B, the most similar sample is the other replicate at that time point. Comparing replicates A and B with replicate C, we see a discordance in two stages during the late tailbud period (*Figure S2.A,B*). We also computed the PCC for all replicate protein and phospho-form trends compared to a scrambled background (*Figure S2.C*). The higher reproducibility of the protein data is consistent with the fact that we are measuring multiple peptides per protein, whereas the increased noise for the phospho-forms is consistent with measuring only a single peptide. All three replicate phospho-form and associated protein measurements for phosphorylation T-429 on the spliceosome protein SF3B1 are shown in Figure 1.C. The median PCC for these phospho-form replicates is the same as the median PCC for the distribution of this metric for all measured phospho-form replicate pairs. The median PCC for the SF3B1 relative protein measurements is lower than the median PCC for the distribution of this metric for all measured protein replicate pairs (0.9).

### The most frequent protein trajectories map to the expected progression of developmental processes

To identify shared trajectories of protein change through development, we clustered the median trends of all proteins using k-means clustering (*Figure 2.A*; See *Figure S3* for the original clusters). The first two clusters are the flattest, and the following six clusters are ordered by the timing of their most prominent temporal event. Fifty-five percent of proteins are in the flat clusters (Clusters 1 and 2), consistent with previous findings from our laboratory [7]. Cluster 3 represents the very small number of proteins that decrease precipitously at oocyte maturation (Stage VI oocyte to egg). Cluster 4 represents the slightly larger number that decrease during the period of cleavage following fertilization. Twenty-one percent of proteins are in cluster 5, which increases gradually following fertilization and remains high until late tailbud (Stage 30), when it slowly declines. Clusters 6, 7, and 8 all increase dramatically no earlier than after the last tailbud stage (Stage 30).

**Figure 2.**
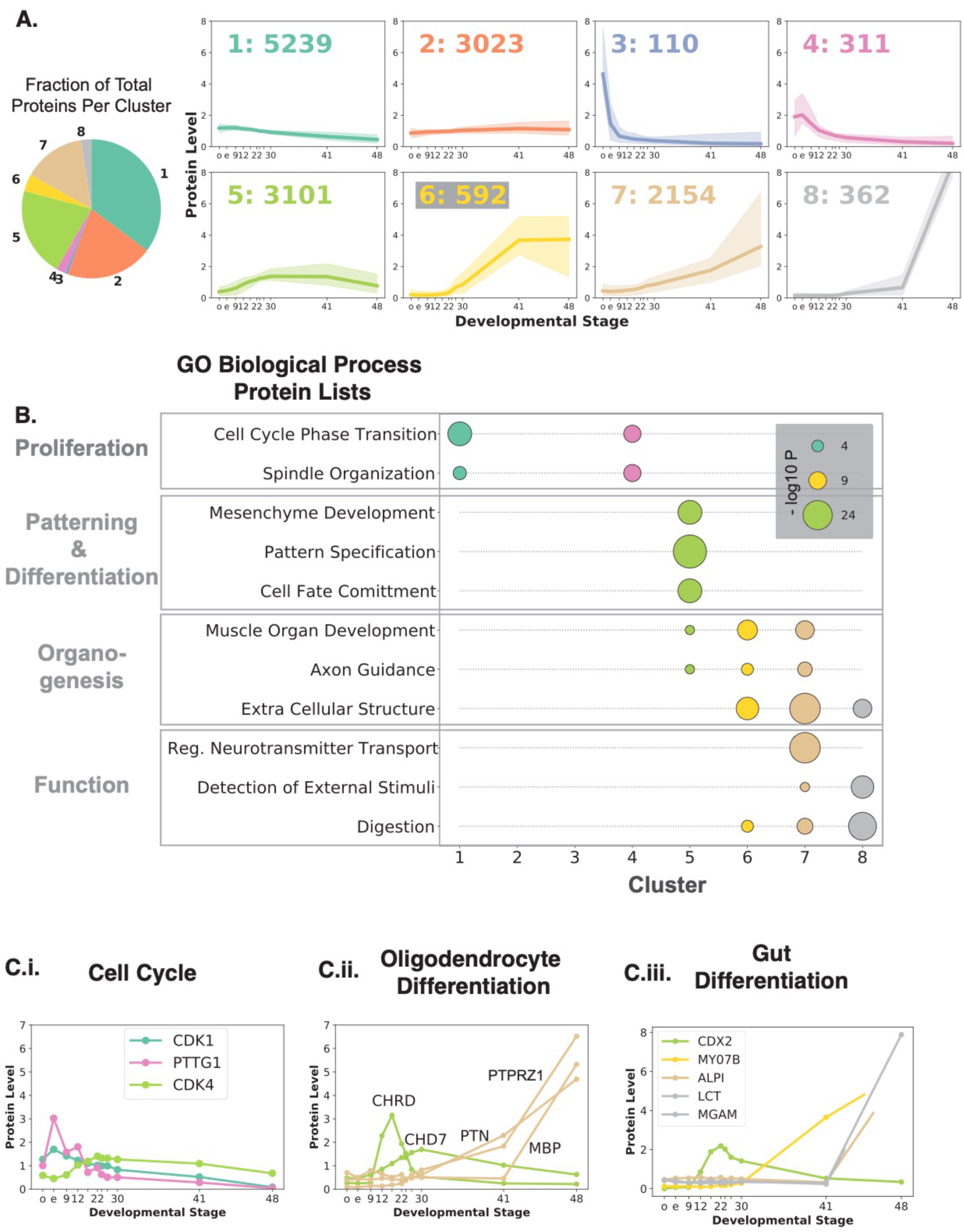
Distinct Protein Expression Clusters Correspond to Major Developmental Transitions. **A.** Twelve k-means clusters of relative protein trends were merged into eight summary trends ordered by the time that the first dynamic event in the trend occurs. The median trend of the cluster is the colored line; the ten to ninety percentile region is shown with shading. The pie chart shows the fractions of total proteins in each cluster. **B.** Over-representation analysis of biological process GO gene sets shows that as development progresses dynamic clusters are associated with proliferation, patterning and differentiation, organogenesis, and tissue function. For each GO protein list shown in black, the p-value for the over-representation of those proteins in the gene set relative to all proteins is shown with a circle, whose area is proportional to the -log10 p-value (hypergeometric) and has the appropriate cluster color. All GO protein lists are overrepresented in at least one cluster with FDR <= 5%. (See *Spreadsheet S1* for the entire set of over-represented sets). **C. (i)** Median trend relative protein levels consistent with the transitions in the mitotic cell cycle across development. Securin/PTTG1 is highest in the metaphase arrested egg and decreases following fertilization. CDK4 levels increase following Stage 9 as the embryo enters the period where entry into S phase is under regulation. (See Figure S4 for individual replicate trends). **(ii,iii)** Expression of proteins connected to tissue differentiation and organogenesis: for oligodendrocytes **(ii)** and gut **(iii)** (See Figure S4 for individual replicate trends). The plots illustrate the temporal progression through these phases, as supported by the GO overrepresentation analysis.

We next analyzed the clusters using Gene Ontology (GO). Selected gene sets that are overrepresented in the clusters that best represent known developmental processes are illustrated in *Figure 2.B* (*Methods; Spreadsheet S1* for all overrepresented GO sets). Cell cycle proteins were notably overrepresented in cluster 4. The precise synchrony of early stages makes it easy to measure these cell cycle dependent changes at the protein level. Several of the cyclin proteins -- Cyclins B1, B2, E1, A1 -- are in cluster 4, where the levels drop as the cell cycle slows down after the mid-blastula transition (Developmental Stage 8) and through gastrulation [18,19]. The same is true for other cell cycle proteins whose levels oscillate during the cell cycle, such as Securin (PTTG1, *Figure 2.C.i., Figure S4.A.*) [20]. The decrease in mitotic protein levels after the egg in cluster 4 is consistent with both the loss of synchrony and the decrease in the mitotic index. Mitotic proliferation related gene sets are not overrepresented in any of the later clusters; however, proteins involved in regulation of the G1/S transition, such as Cdk4, cyclin D, and RB1, are found in cluster 5, suggesting that regulated cell proliferation continues at a high level throughout development. The gene sets “cell cycle phase transition” and “spindle organization” are also overrepresented in the first flat cluster, including the master cell cycle kinase CDK1, the CDK1 regulator ENSA, the anaphase promoting complex protein CDC16, and the aurora kinase regulator BORA.

After the mid-blastula transition (MBT) which begins after the 12^th^ cell division, the cell cycle slows down, transcription begins, and a program of differentiation ensues. “Pattern specification” and “cell fate commitment” gene sets are both overrepresented in cluster 5. The gene sets associated with organogenesis and specific cellular functions (“function”) are almost all overrepresented in more than one cluster. “Muscle organ development” and “axon guidance” are overrepresented in all of the clusters that increase after fertilization except for the cluster that increases after the early tadpole stage. This is consistent with the fact that by Stage 41 the tadpole musculature has been established and the development of axons and nerve fibers is extensive [21]. The gene sets related to organ function, such as “regulation of neurotransmitter transport,” “detection of external stimuli,” and “digestion,” are all overrepresented in at least one of the last two clusters, which increase in level from the tadpole at Stage 41 to the true juvenile at Stage 48. This is the period when we expect specific cellular functions to emerge in organs. Examples of the temporal patterns of specific proteins of oligodendrocyte differentiation (*Figure 2.C.ii, Figure S4.B.*) and gut differentiation (*Figure 2.C.iii, Figure S4.C.*), as well as their precursors, exemplify progression of differentiation leading to organ function. For example, the neural lineage progenitors including glial cells are physically located in the dorsal part of the embryo. Chordin (CHRD) is a secreted protein in the Mangold-Spemann organizer that patterns the dorsal ventral axis during gastrulation and neurulation, particularly the brain and spinal cord [22]. CHD7 is a bromodomain ATP dependent remodeling protein that stabilizes gene expression and is involved in specifying the oligodendrocyte lineage [23]. PTPRZ1 is a receptor protein tyrosine phosphatase that inhibits embryonic oligodendrocyte differentiation in mice, but its inhibitory function is blocked by the ligand pleiotrophin (PTN) [24]. Myelin basic protein (MBP) is one of the most abundant proteins in myelinated sheaths, which are the terminal differentiated product of the oligodendrocyte lineage. Protein levels of MBP increase only after Stage 41. This is consistent with mRNA levels of the brain isoforms of MBP not being detectable at Stage 40, and MBP protein being detectable in the Stage 47 spinal cord [25].

In contrast to neural progenitors, which develops on the dorsal side, the gut is a ventral organ. The transcription factor CDX2 is involved in regulation of the posterior gut epithelium [26,27] and persists in the gut stem cell population. A histological marker, MYO7B, is an unconventional myosin expressed in microvilli of enterocytes in the posterior gut in mice (and also in the kidney) [28]. Intestinal type alkaline phosphatase (ALPI), lactase-phlorizin hydrolase (LCT), and intestinal maltase-glucoamylase (MGAM) are all digestive enzymes found in the small intestine. These proteins would be necessary for intestinal function. Food is first found in the gut at Stage 46 [21], suggesting that the gut is well differentiated at that stage. Thus, the measured protein dynamics are consistent with the progression from patterning and fate specification (CHRD, CHD7, CDX2) at the transcriptional level to regulated commitment and emergence of structure (PTPRZ1, PTN, MYO7B), to juvenile function (MBP, ALPI, LCT, MGAM). We expect that specific proteins that follow a similar trajectory in other lineages could be found in the data and may be identified by a combination of single cell transcriptional analysis and proteomics.

### Major changes in the proteome during organogenesis

We can subdivide the proteomic dataset into two periods: proteins involved in proliferation, patterning, and specification in the first period; and those acting in organogenesis and function in the second period. Based on the median trends of the clusters in Figure 2, we chose Stage 30 as the breakpoint between the two periods. For each protein in the database we can ask what fraction of the overall magnitude of protein level change happens before Stage 30 as compared to the overall change from the oocyte to the feeding tadpole, Stage 48 (*Figure 3.A*; *Figure S5*). The ratio of these two values, which we call the fractional fold change (FFC), will be greater than or less than 0.5 if the majority of the change is achieved before or after Stage 30. Figure 3.A illustrates these metrics for the nuclear RNA binding protein HNRNPU and the complement pathway receptor CD55. The FFC value of HNRNPU is very close to one, whereas that of CD55 is 0.18. Thus, for HNRNPU more change occurs before Stage 30, while the opposite is true for CD55.

**Figure 3.**
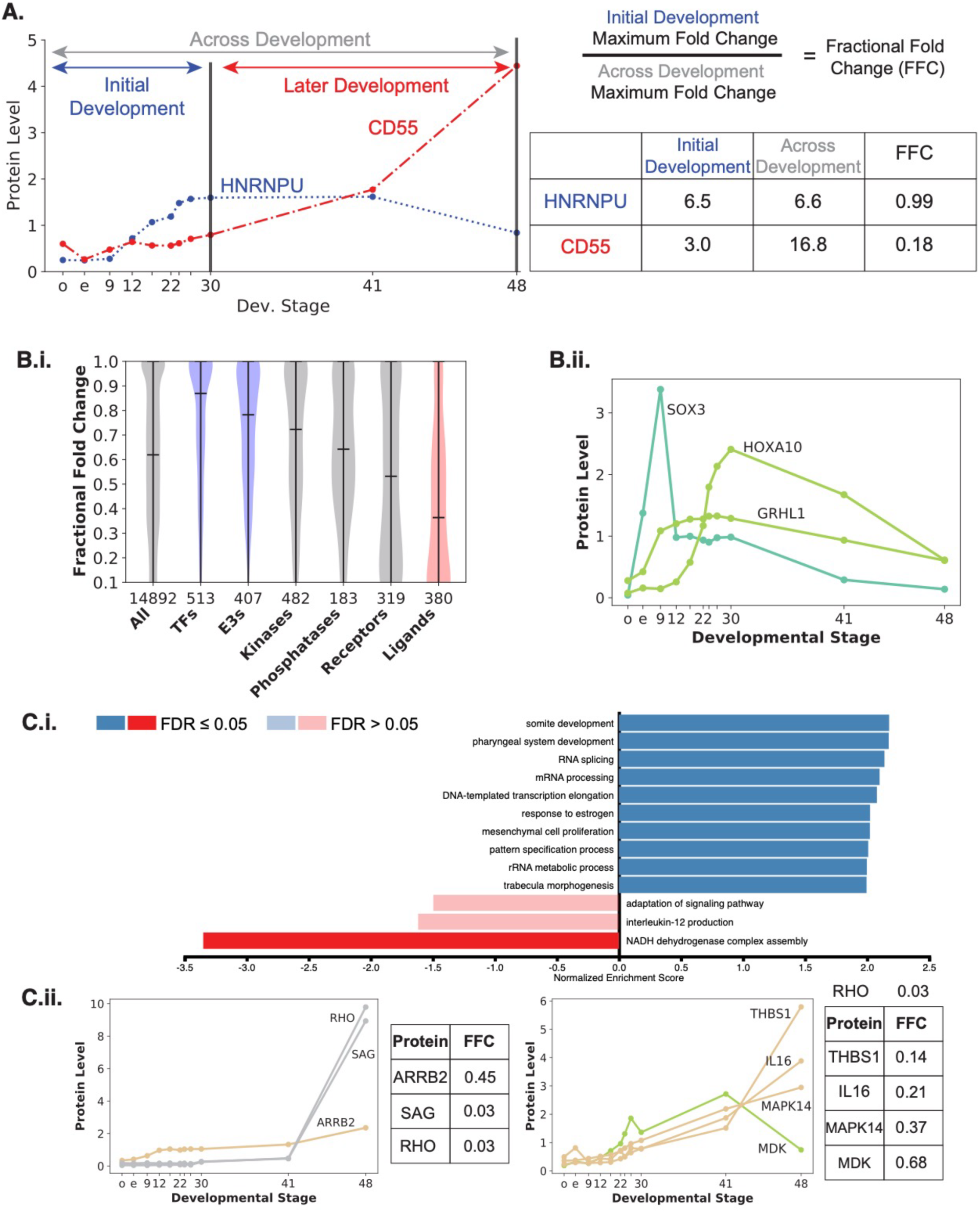
Gene Expression Regulators Show Pre-Stage 30 Dynamics While Cell Communication Proteins Exhibit Post-Stage 30 Changes. **A.** Introducing the fraction fold change metric (FFC). The measured time period is split at Stage 30 into “initial development” (blue) and “later development” (red). In order to determine how much of the maximum fold change across the entirety of development occurs during “initial development,” we divide the maximum fold change during initial development by the maximum fold change across development. We illustrate the metric with the two examples from the data – HNRNPU and CD55 – that have very different fraction fold changes. (For further visualization see *Figure S5*.) **B. (i)** Violin plots of the median trend FFCs for the six different classes and all proteins (center black line = median; top and bottom black lines = max and min, respectively). The medians of transcription factors (TFs), E3 ubiquitin ligases (E3s), and ligands differ significantly from all proteins for the median trend and all three replicates (p < 0.001; Kruskal Wallis test, Bonferroni critical value for multiple comparisons with post-hoc Dunn’s test). Violin plots for the individual replicates are shown in *Figure S6*. **(ii)** Examples of transcription factor relative trends with FFCs equal to one. The data shown is the median trend of the replicates for which the protein is measured. The trend lines are colored according to the clusters in Figure 2.A. Individual replicate trends are shown in *Figure S7*. **C. (i)** Gene Set Enrichment Analysis (GSEA) results for the median trend FFC metric using Biological Process GO sets shows that processes related to gene expression and cell fate differentiation are enriched for FFCs closer to 1 compared to the background of all measured proteins. In contrast, processes related to cell-environment signaling and the immune response are depleted for FFCs closer to 1. **(ii)**Median protein trends for some well-known members of the (left) “adaptation of signaling pathway” and (right) “interleukin-12 production” gene sets. Individual replicates are show in *Figure S8*.

We calculated the FFC for different classes of regulatory proteins: transcription factors (TFs), ubiquitin ligases, kinases, phosphatases, receptors, and ligands [29] (*Figure 3.B.i, Figure S6*). The median FFC for TFs, ubiquitin ligases, kinases, and ligands are all significantly different from the median of all proteins. The median FFCs of TFs and ubiquitin ligases are larger than the median of all proteins (0.62) and higher than 0.5 (0.87 and 0.78, respectively). This indicates that TFs and ubiquitin ligases regulate gene expression and patterning events before Stage 30, and with ubiquitin ligases also regulating progression through the cell cycle during the period of high mitotic index early in embryogenesis [30–32] [33,34]. As examples, the transcription factors SOX3, GRHL1, and HOXA10 are required for early neural specification, epithelial development, and anterior-posterior organization/myeloid differentiation, respectively, all have FFCs equal one (*Figure 3.B.ii, Figure S7*). In striking contrast to TFs and E3s, ligands have a median FFC that is smaller than that of all proteins (0.36, respectively, as compared to 0.62), suggesting that they are generally expressed locally in tissues at the time of the tissue function.

To assess what other biological processes are associated with the periods before, as opposed to after, Stage 30, we performed Gene Set Enrichment Analysis using the FFC as our quantitative metric. Multiple GO biological process gene sets related to transcription, mRNA processing and metabolism, and ribosomal RNA transcription (“rRNA metabolic process I”) are statistically significantly enriched for FFCs closer to one compared to the set of all measured proteins (*Figure 3.C.*). This is consistent with the distribution of transcription factor FFCs and the obvious importance of regulation of gene expression for patterning and differentiation early in development. It is also known that transcription of ribosomal rRNA begins during gastrulation (before Stage 30) [35].

The GO gene sets that have FFCs closer to zero (changes late in development) include “adaptation of signaling pathway” and “interleukin-12 production.” The adaptation of signaling pathway includes proteins involved in response to light stimulation (e.g. RHO and SAG) as well G-protein coupled signal proteins (e.g. SAG and ARRB2) (Figure 3.C.ii, *Figure S8*). Interleukin-12 is a cytokine that has multiple roles, including in the differentiation of T-cells. The “interleukin-12 production” pathway includes ligands THBS1, IL16, and MDK, as well as kinase MAPK14 (Figure 3.C.ii, *Figure S8*). At the cellular level these proteins are involved in intracellular/extracellular communication and, on the organismal scale, with interaction of the embryo with its environment. This is consistent with the basic difference between the Stage 30 tailbud embryo and Stage 48 tadpole -- a difference not in patterning or specification, but in the capacity to interact with the external environment (sensory organs) and particularly to interact with the internal environment, such as required for a functional digestive system, nervous system, and immune system. By Stage 30 physiological function has replaced establishment of organization as the preoccupation of the organism.

### Embryonic growth: quantitative changes in non-yolk protein between the late tailbud and juvenile tadpole

Up to this point we have considered the relative, rather than the absolute changes of proteins across development. However, by combining the relative protein changes with the changes in mass of total non-yolk protein we can calculate the absolute protein abundances across development. The total amount of protein per embryo (average of 10 embryos) after removal of yolk for all of the timepoints in our proteomics dataset except for the oocyte is shown in *Figure 4.A* (*Methods*). Yolk, having been endocytosed from protein produced in the mother’s liver, is held until neurulation before it begins to break down [10]. In the egg, 85% of protein is yolk (*Figure S9*). By Stage 48, there is no detectable yolk protein. In early embryogenesis before Stage 30, there is no measurable increase in non-yolk protein amount in the embryo. Later, the average amount of non-yolk protein increases 1.3-fold between Stage 30 and Stage 41, and 1.5-fold between Stage 41 and Stage 48, for an overall two-fold change from Stage 30 to Stage 48. If all the yolk protein were converted into amino acids, and if all the amino acids were converted into new protein, we would expect protein content to increase about 7.5-fold, not 2-fold. One explanation for this discrepancy could be that the amino acids derived from yolk are mostly metabolized as an energy source. This would be consistent with the finding in the amphibian *Rana fusca* that excretion of nitrogen increases 10-fold at the time of embryo hatching [36,37].

**Figure 4.**
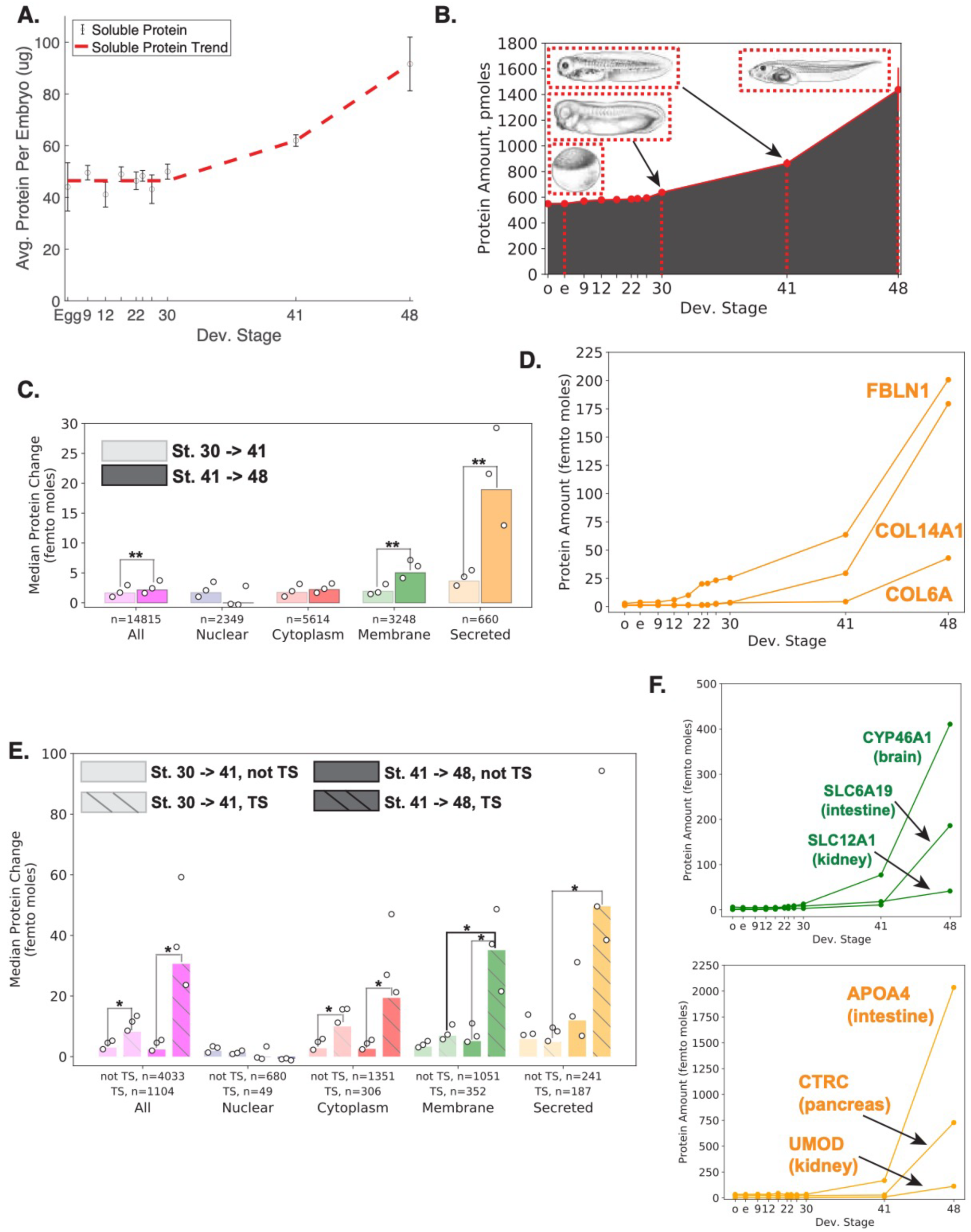
Two-Fold Proteome Mass Increase from Late Tailbud to Tadpole is Predominantly Driven by Tissue-Specific Proteins. **A.** Amount of protein per egg/embryo (average of ten) at ten developmental stages. The mean of three technical replicates is shown with 95% confidence intervals. **B.** The total amount of protein, in picomoles, at the measured developmental stages. The dots are the mean of three replicates and the error bars are the standard deviation. Red lines show the developmental stages of embryo drawings. **C.** Median trend (bars) absolute change of proteins by annotated localization for Stage 30 to 41 (lighter shade) and Stages 41 to 48 (darker shade). Open circles are the values for each replicate for each subset. Statistical significance was determined using the Kruskal Wallis test with Bonferroni correction for 20 comparisons (α = 0.0025). **D.** Absolute protein amount median trends for three different extracellular matrix proteins. Individual replicates are shown in *Supplemental* Figure 11*.A*. **E.** Median trend (bars) absolute change of proteins with the localization classifications additionally stratified by tissue specificity (TS) (not tissue specific: no black lines, tissue specific: black lines; left and right, respectively) for Stages 30 to 41 (lighter shade) and Stages 41 to 48 (darker shade)). Open circles are the values for each replicate for each subset. Asterisks show the comparisons where the median trend and all three replicates are statistically significantly different, where statistical significance for a comparison is assessed with the Kruskal Wallis test, Bonferroni correction for eighty comparisons (α = 0.000625). **F.** The absolute amount of protein across development for three tissue specific membrane proteins and tissue specific secreted proteins. Individual replicates are shown in Supplemental Figure 11.B,C.

To convert the relative changes in protein to the molar amount of protein change for the proteins in our dataset, we cross-referenced our data with the reference of absolute protein concentrations in the egg to determine the relationship between mass spectrometry signal and protein concentration [2] (*Methods; Figure S10*). Using these measurements, we are able to determine the number of moles for each protein in our dataset and the change in the total amount of moles of protein across our time series for the measured portion of the proteome (*Figure 4.B*.; *Methods*). This method of calculation shows a 2.3-fold change, slightly greater than the mass measurement.

### Changes between early and late organogenesis

The amount of protein increase between Stages 30 and 41 is not very different from the amount of protein increase between Stages 41 and 48. We wanted to know whether the composition of protein changes during these periods is also similar. Our analysis of regulatory protein classes revealed that receptors and ligands are more dynamic after Stage 30 than for the set of all measured proteins. Encouraged by this finding, and by the desire to include larger numbers of proteins in our analysis, we decided asked about the intracellular distribution of proteins. We classified 75% of proteins as either nuclear, cytoplasmic, membrane, or secreted (*Methods*). Between Stages 41 and 48, the median change of membrane and secreted proteins is larger than that for all proteins, and the median change between Stages 41 and 48 is larger than that between Stages 30 and 41 (*Figure 4.C*). Examples of secreted proteins that show large molar increases between Stages 41 and 48 are extracellular matrix proteins fibulin-1 (FBLN1), collagen alpha-1(XIV) chain, and collagen alpha-3(VI) chain (*Figure 4.D, Figure S11*). In contrast to membrane and secreted proteins, the median change of nuclear proteins is much smaller between Stages 41 and 48 than between Stages 30 and 41. The median increase for cytoplasmic proteins is very similar. Figure 4.C shows the median change within a class, but we can also sum the changes to determine the contributions to the total protein mass of the different classes. Between Stages 30 and 41, membrane and secreted proteins constitute 34% of the protein increase; between Stages 41 and 48, they constitute 52% (considering only those proteins that have a localization class). Thus, despite the similarity in change of total protein, the composition of change is different for the two periods.

The prominent changes observed in later development likely reflect development of multiple organs and their constituent cell types, which initiate their physiological functions between Stages 41 and 48. We explored how the increase in protein levels relates to tissue specificity. In this broad characterization, we classified proteins as non-tissue-specific if they were either present in the egg or “not tissue enriched” in adult organs. We classified proteins as tissue-specific if they were both not present in the egg and “tissue enriched” in adult organs [38] (*Methods*). By these criteria, 7.5% of proteins in the dataset are tissue specific. Despite their minor representation, tissue-specific proteins constitute 37% of the total molar increase between Stages 41 and 48. The set of membrane proteins that are tissue-specific increases more than non-specific membrane ones between Stages 41 and 48 and more between Stages 41 and 48 than between Stages 30 and 41 (*Figure 4.E*). Two common types of tissue-specific membrane proteins that increase after Stage 30 are cytochrome P450 monooxygenase enzymes (e.g. CYP46A1 in the brain) and solute transporters (e.g. SLC12A1 in the kidney and SLC6A19 in the intestine) (*Figure 4.F, Figure S11*). The set of secreted proteins that are tissue-specific increases more between Stages 41 to 48 than between Stages 30 to 41 (*Figure 4.E*). Examples of tissue-specific secreted proteins that show a particularly dramatic increase after Stage 41 include apolipoprotein A-IV (APOA4) in the intestine, chymotrypsin-C (CTRC) in the pancreas, and uromodulin (UMOD) in the kidney (*Figure 4.F, Figure S11*). Cytoplasmic tissue-specific proteins also increase more than non-tissue-specific cytoplasmic proteins during both of these periods. Many of the tissue-specific cytoplasmic proteins are expressed in muscle tissues. Trends for two tissue-specific cytoplasmic muscle proteins, myomesin-2 (MYOM2) and myozenin-2 (MYOZ2), are shown in *Figure S12*. The large molar increase of tissue-specific proteins is consistent with the overall increase in proteins from the egg stage (*Fig 4.A)*, where these proteins are either not present at all or at very low levels relative to the levels in the juvenile tadpole. Though the larger median increase of tissue-specific proteins than other proteins is a feature of the entire period after Stage 30, their contribution to the increasing proteome is more significant after Stage 41. Taken together, these observations suggest that a dramatic change in the biochemistry of protein production occurs after Stage 30, resulting in an increase of the soluble proteome by mass, and that a second transition event after Stage 41 results in a change in the composition of the types of proteins that are made during the crucial period when organs initiate their physiological function.

### Changes in protein phosphorylation are to proliferation and control of gene expression

In our efforts to understand the role of protein phosphorylation in development, we co-analyzed the phospho-form and corresponding protein abundance for every phospho-form. Simultaneous protein and phosphorylation analysis distinguishes between two scenarios: phosphorylation sites that are inherently dynamic, and constitutive sites that only appear to change simply because their underlying protein levels change proportionally, as shown in these schematics in *Figure 5.A.* When protein and phosphorylation patterns are not coincident, this indicates a change in the proportion of phosphorylated protein, suggesting the phosphorylation likely serves a regulatory purpose (*Figure 5.A.iii*). To distinguish quantitatively the dynamical nature of the phospho-forms, we calculate the Pearson Correlation Coefficient of the phospho-form and protein trends, which we term “PP Corr” (*Figure S13.A*). Dynamic phospho-forms will have a low PP Corr; however, a low PP Corr can also occur when neither protein nor phospho-form levels change. In such cases the discordance is assumed to be due to noise. To distinguish these possibilities, we also determine the maximum fold change of the phosphorylation trend across the time series, which we term “Phos FC” (*Figure S13.B*). We used these metrics to compare k-means co-clusters of the protein and phosphorylation trends (*Figure 5.B., Figure S14* for the original twelve co-clusters). (These co-clusters are labelled A-G to distinguish them from the protein-only clusters in *Figure 2.*) The median values of PP Corr and Phos FC for each cluster (with bars covering the interquartile – 25^th^ to 75^th^ percentile -- region) are shown in *Figure 5.C.* along with some of the biological process GO sets that are overrepresented (See *Spreadsheet S2* for complete lists).

**Figure 5.**
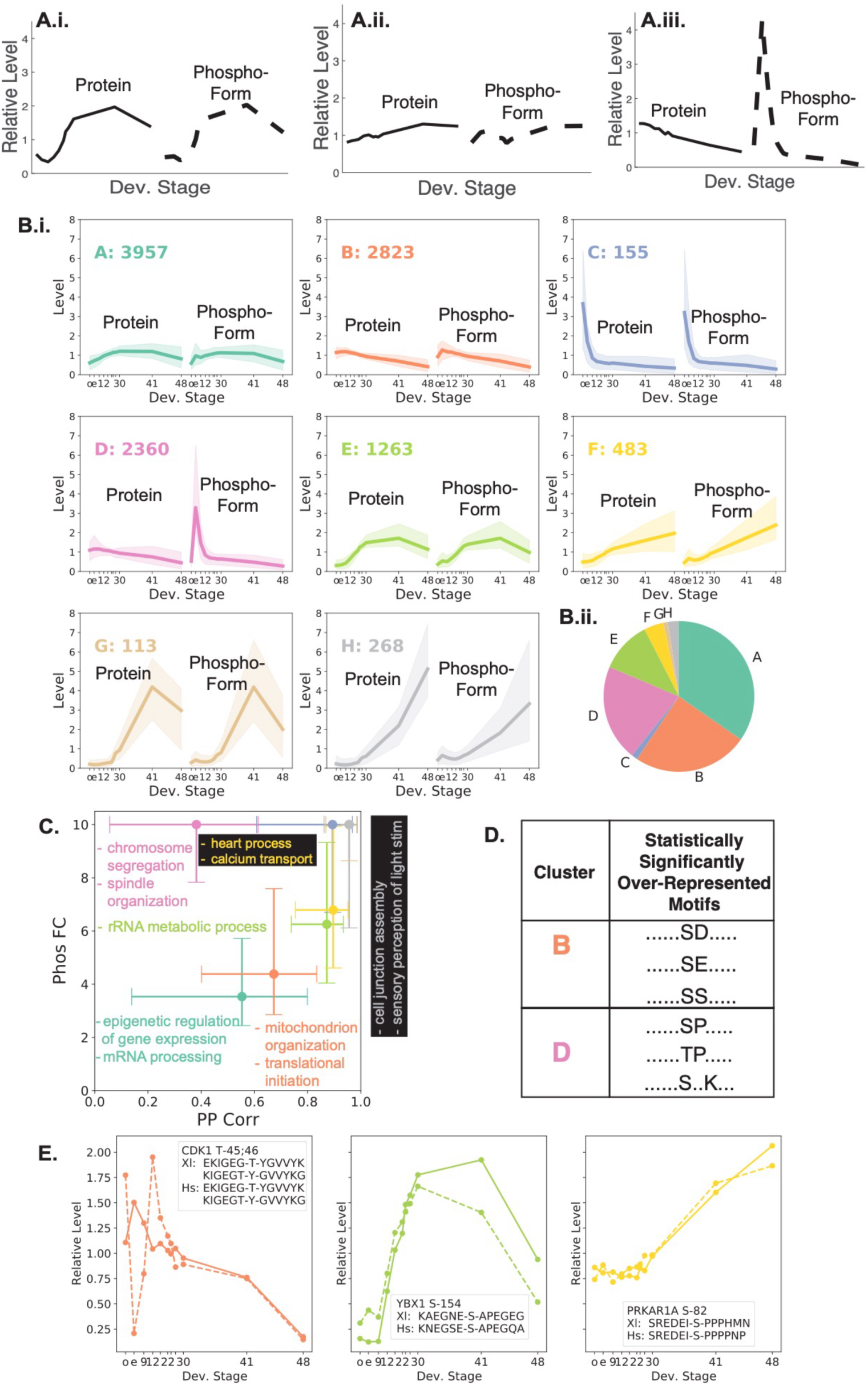
Mitotic cell cycle and gene expression related phosphorylation dominate the phospho-proteome. **A.** Phospho-form dynamics are investigated together with their underlying protein dynamics and without normalization to the protein trend in order to distinguish **(i)** phospho-forms that are similar to their protein trend but have significant changes in magnitude from **(ii)** those that are also similar but do not change very much during development. **(iii)** Other phospho-forms have very different trends than the protein they are measured on. **B.** Over 50% of phospho-forms are relatively flat and the majority of dynamic phospho-forms show most of their increase before the tadpole stages. **(i)** The protein and phospho-form trends were clustered together as a vector of twenty-two timepoints. Twelve original *(Supplemental* Figure 14) k-means clusters were merged into eight summary clusters. Cluster medians (dark colored lines) and ten to ninety percentile regions (lighter colored area) are plotted separately for proteins and phospho-forms to aid interpretation. **(ii)** The fraction of the total phospho-forms that are in each cluster is represented by a pie chart where the colors are the same as the cluster medians. **C.** Cluster median Pearson correlation between phospho-form and protein (PP Corr) and phospho-form maximum fold change across development (Phos FC) are shown for all clusters with error bars representing the 25 to 75% interquartile range the same co-cluster colors. The maximum allowable phospho-form max. fold change across development is 10. GO Biological Process protein sets that are overrepresented for a given cluster with FDR <= 5% are shown in the same cluster color. See *Spreadsheet S2* for the complete set of overrepresented sets. **D.** Cluster B phospho-forms are statistically significantly overrepresented for multiple acidic motifs, and cluster D phospho-forms are statistically significantly overrepresented for multiple proline directed motifs. See *Spreadsheet S3* for the complete lists of statistically significantly overrepresented motifs. **E.** Examples of phosphorylations from three different co-clusters that have a homologous human phosphorylation. The median trend phospho-form (dashed) and protein (solid) lines are shown where the color matches the phospho-form’s co-cluster. Individual replicate data is shown in Supplemental Figure 17.

Together, co-clusters A and B, comprising 59%, of all the data show parallel changes in the protein and phosphorylation abundance. They have median Phos FCs of 3.53 and 4.38, respectively, and median PPCorrs of 0.55 and 0.67, respectively. Co-clusters D and E are 21% and 11% of the phosho-form data, respectively. With median PhosFCs of 10 and 6.25, their phosphorylation changes are larger than those of co-clusters A and B. GO sets related to gene expression are overrepresented in both clusters A and E. In both cases the phosphorylation levels decrease between Stages 41 and 48, which is in agreement with the dynamics of gene expression proteins across development and suggests that the majority of phosphorylation related to gene expression is constitutive. Cluster D has the smallest median PP Corr, suggesting phospho-forms that are regulatory. Gene sets “chromosome segregation” and “spindle organization” are overrepresented in Cluster D, which is consistent with the fact that the phosphorylation dynamics match the varying roles of the cell cycle during development. The small PP Corr is consistent with the importance of phosphorylation for specific regulation of cell cycle progression [39] and the role of phosphatases in removing mitotic phosphorylation, which must result in divergence of protein and phosphorylation dynamics [40,41]. Cluster F is the largest cluster (4.2% of the data) where in contrast to co-clusters A through E, the median phosphorylation abundance increases after Stage 30. Co-clusters C, G, and H together comprise just under 5% of the data but represent some of the most dramatic changes (all have median PhosFCs of 10).

Fluctuations in phosphorylation levels arise from altered kinase/phosphatase activity or, in certain cases, differential protein degradation rates. To connect the protein substrate phosphorylations to the kinases responsible, we used two approaches. We first determined the phosphorylation motifs that are overrepresented in the clusters [42–44] (*Figure 5.D.; Methods; Supplemental Table 1* for complete lists). We then took advantage of annotation of specific phosphorylations from human datasets, and we developed a custom pipeline that uses global and local sequence homology to identify human phosphorylations that are homologous to our measured Xenopus ones (See *Figure S15* and *Methods)*. We leveraged kinase-substrate motif data from well-characterized pairs using the Phosphosite/Kinase module within the Webgestalt analysis suite [45].

The acidic motifs SD and SE are overrepresented in Cluster B. These motifs are related to the Casein Kinase 2 motif [46]. The CK2 group has the largest overlap with the human mapped phosphorylations from Cluster B that include these motifs *(Methods, Spreadsheet S3)*. The CK2 kinases are constitutively active, which is consistent with the similarity of the protein and phospho-form profiles in this cluster. Examples of phosphorylations in this set in Cluster B are shown in Figure S16. CK2 is known to phosphorylate proteins involved in ribosomal RNA processing and ribosome biogenesis, processes that are overrepresented in co-cluster E [47,48]. Examples of nucleolar proteins with phosphorylations in acidic motifs are mRNA protein turnover protein 4 homolog (MTRO4) and ribosome biogenesis protein BMS1 homolog (BMS1), which function in pre-60S subunit assembly and pre-40S subunit assembly, respectively (*Figure 6D, left*) [49,50]. When the entire dataset is considered together, phosphorylations in acidic regions have the highest PP Corr compared to other broad motif classifications (see below and Figure 6); this is consistent with known properties of CK2 and evidence for low phosphatase selectivity for phosphorylations proximal to acidic residues [51,52].

**Figure 6.**
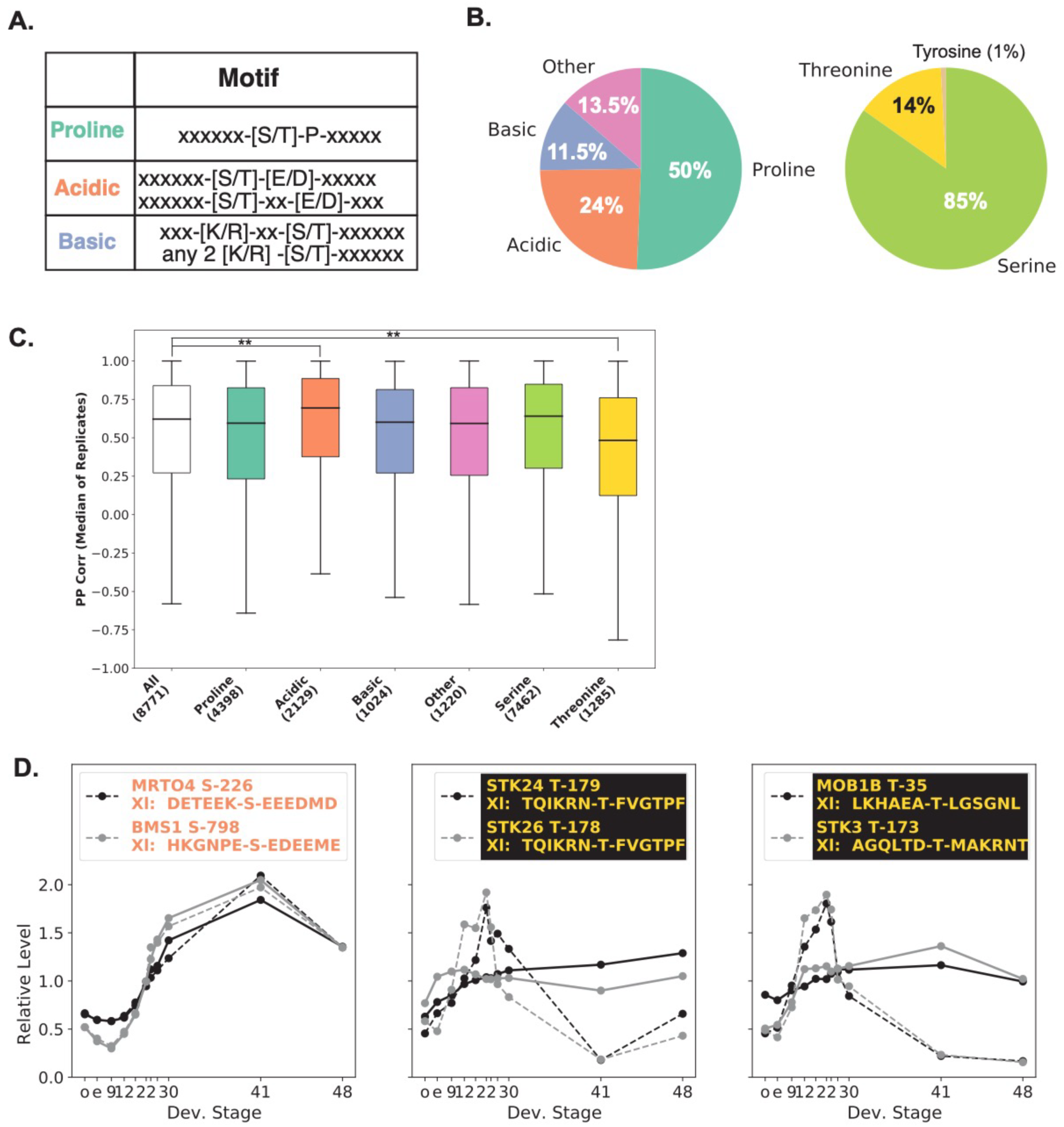
Acidic Motif Phosphorylations Correlate Strongly with Protein Trends, Unlike Threonine Phosphorylations. **A.** Definitions of the motif classifications. **B.** Fraction of phospho-forms by **(left)** motif classification and **(right)** phospho-acceptor residue. 50% of phospho-forms have proline-directed motifs. The vast majority of phospho-forms have serine phospho-acceptor residues. **C.** Phospho-forms with acidic motifs are more correlated with their associated protein trend than the set of all phospho-forms, and threonine phospho-acceptor phospho-forms are the least correlated (** = p < 0.001; Kruskal Wallis test, Bonferroni critical value for multiple comparisons including comparisons of the median values and the three replicates). See *Supplemental* Figure 18 for the same visualization for the three replicates separately. **D.** Examples of phosphorylations (dashed lines) and associated protein (solid lines) with high PP Corr values (acidic motifs, left) and lower PP Corr values (threonine phosphorylations, center and right). The threonine phosphorylations examples increase in phosphorylation level after the midblastula transition (just before Dev. Stage 9). Individual replicate data is shown in Supplemental Figure 20.

The proline directed motifs SP and TP are overrepresented in Cluster D. These proline-directed motifs are associated with multiple kinases including cyclin-dependent kinases, most prominently CDC2/CDK1 [46] (*Methods, Spreadsheet S3* for complete list). This is consistent with high activity of this master-cell cycle kinase in the egg (Figure 5.E.left, Figure S17.A; as expected the inhibitory doubly phosphorylated T-14 and Y-15 (in human) phospho-form of CDK1 is at its lowest level (least inhibited) in the egg). The remaining panels of Figure 5.E (Figure S17) show examples of phosphorylations in Clusters E and F where the measured Xenopus phospho-form maps to a known human phospho-site. The relative phosphorylation dataset includes the complete human mapping results.

### Threonine phosphorylations show weaker correlation between protein and phospho-form dynamics

A surprising finding from our clustering approach is that the only cluster with a PP Corr less than 0.5 is the mitotic phosphorylation cluster. However, the large interquartile regions of many of the clusters show that the data also include phospho-forms with trends divergent from that of their proteins that do not peak at the egg. We decided to classify the phospho-form data instead by the identities/properties of the phosphorylated amino acid and the surrounding (+/- six) amino acids. We used a published motif classification scheme that classifies motifs with the following hierarchy: proline-directed, acidic, or finally basic [53]. Phospho-forms with motifs not meeting any of these classifications were grouped as “other” (*Figure 6.A; 6.B.i*). We also grouped our phospho-forms by phospho-acceptor residue (serine vs. threonine) (*Figure 6.B.ii*). The preponderance of serine phospho-acceptors over threonine is consistent with previous studies in mouse tissues and human cell lines which reported 11-15.5% of phosphorylation on threonine [53–55]. The threonine phospho-forms have the lowest median PP Corr of all classes (*Figure 6.C, Figure S18*). There is evidence from multiple studies that this lower PP Corr may be attributable, not to the kinase regulation, but to phosphatase regulation [52,56,57]. An alternative possibility could be that threonine phospho-acceptor phospho-forms appear less correlated to protein levels because of noise. In fact, the median for the distribution of Phos FC median values for threonine sites is larger than that for serine sites, suggesting that the low PP Corr threonine are due to dynamic phosphorylation regulation, rather than noise (Figure S19). Many of the threonine phospho-acceptor phospho-forms are proline directed (63%). This is consistent with the overrepresentation of the TP motif in the mitotic phosphorylation cluster that has the lowest PP Corr, and is also consistent with studies of phosphatase regulation of mitotic exit [40,52,56,57].

There are threonine phospho-acceptor phospho-forms with lower PP Corrs that increase in phosphorylation level after the MBT, in contrast to the pattern of high mitotic phosphorylation in the egg. Figure 6.D.center *(Figure S20)* shows examples of threonine phospho-forms on the kinases STK24/MST3 and STE26/MST4. For MST3, the phospho-form is at the active site, and the phosphorylation induces activity [58]. STK24 and STK26 regulate the cytoskeleton, cell polarity, and Golgi Apparatus positioning [59–61]. Figure 6.D right *(Figure S20)* shows threonine phosphorylations on the Hippo signaling pathway proteins STK3/MST2 and MOB1B. MOB1B is a mediator between the upstream kinases STK3/MST2 and MST1 and the downstream kinases, which include LATS1 [62]. The T-35 phosphorylation on MOB1B is a substrate phosphorylation of STK3 [63]. The striking accumulations of these phosphorylations on the Hippo machinery during the neurula and early tailbud period, and their decrease, thereafter, suggests a very specific developmental role for YAP signaling at this stage of embryonic development. This may represent a concerted transition in the whole embryo, like the MBT, or it could just be a very strong signal confined to a small set of cells.

## DISCUSSION

We have produced novel and extensive datasets that provide information about protein and phosphorylation changes of interest during embryonic development. The synchrony of the developmental processes and the large size of the Xenopus embryo are both essential for proteomic analysis. The phospho-proteome is even more difficult to examine in most model systems. For this reason, this data, as extensive as it is, is still far from complete, but it offers a substantial look into changes on the proteomic level in an early vertebrate system. Xenopus may be the optimal common developmental model for proteomic and phospho-proteomic analysis at this time, principally because a reasonable number of embryos provide enough material for proteomic analysis, they can be easily synchronized, and because we have deep knowledge of the developmental processes, which in many ways are similar to those in mammals. Because the RNA transcriptome allows much greater depth of coverage, it has been the only practical approach to understanding protein expression. However, in other systems, such as cell culture, we have come to realize that translational control, protein degradation, and protein phosphorylation play an important role in molding phenotype. We are able to report, for the first time in an embryonic system beyond cleavage stage, quantitative protein phosphorylation levels. Due to the present limitations of mass spectrometric methods, we have still only recorded these to a limited depth compared to the expected total phospho-proteome. Likely tens of thousands of phosphorylated residues remain to be measured [11]. The improved depth and temporal resolution during early development did not challenge our previous finding that abundance changes of the majority of proteins are very small across early and mid-stages of embryogenesis. However, by extending the time period of our measurement to the embryo/juvenile transition, we observe multiple biochemical changes of protein and phosphoprotein regulation. There is a two-fold increase in total (non-yolk) protein between Stages 30 and 48. The composition of the proteome shifts after Stage 41 as membrane and secreted proteins increase faster than nuclear proteins. It is not until organ systems become functional that this dramatic change in the proteome occurs.

The most prominent trend of large-scale changes in phosphorylation level are the phospho-forms that are highest in the metaphase-arrested egg. That so many of these phosphorylations are detected, even though the majority decrease dramatically in level by the next time point (Developmental Stage 9), is likely explained by their high occupancy in the mitotically arrested egg. This strongly expected high occupancy at the metaphase arrested egg is due both to the fact that metaphase phosphorylations are present at high occupancy [54,64], and to the fact that the metaphase arrest egg constitutes a perfectly synchronous and homogeneous state. Though the goal of this study was to measure phosphorylations after the midblastula transition, the prominence of cell cycle related phosphorylation in this data demonstrates again the not yet fully tapped power of the Xenopus system for studying cell cycle phosphorylation at the phospho-proteome wide level.

The most prominent protein abundance and phosphorylation changes after fertilization and after the 12 rapid cleavages leading up to the activation of transcription at the midblastula transition were connected with the regulation of gene expression. Because of the ease of measuring nascent transcripts of RNA, most of regulation of, gene expression during development has been considered in terms of transcription factors and temporally and spatially discrete RNA expression patterns. Phosphorylation related to gene expression has mostly been noted for specific well-characterized phosphorylations that regulate transcription factor localization and stability (e.g. beta-catenin in the canonical Wnt signaling pathway). However, the phosphorylation changes that we detected are on proteins expected to be used ubiquitously by the gene expression machinery (e.g. splicing related proteins serine/arginine-rich splicing factor 12 – SRSF12 and pinin -- PNN), suggesting that there is also an embryo-wide level at which the prioritization of regulated gene expression is occurring. Many of these phosphorylations related to gene expression occur in acidic motifs. Phospho-proteome wide studies of phospho-occupancy in yeast and a cell culture model found that the set of phosphorylations in acidic motifs have higher occupancy than the set of all phosphorylations [65,66]. It would be interesting to determine if these phosphorylations are necessary for protein function or are a byproduct of the combination of non-specific phosphorylation by kinases and low phosphatase activity.

The changes in the absolute levels of proteins and the increase of the total non-yolk protein amount after Stage 30 together provide strong evidence that after Stage 30 there are embryo-wide biochemical events necessary for the development of functional organs. We observe large increases in relative protein levels, increase in the total protein in the embryo, and the composition of the proteome shifts to contain a larger fraction of membrane and secreted proteins. Tissue-specific proteins increase dramatically, particularly after Stage 41. In contrast to the change of the proteome, we measure few large changes in the phospho-proteome. Phosphorylated sites that do increase during this period are all on proteins with a concomitant increase and are likely constitutive rather than regulatory in nature. Dramatic change in phosphorylation levels of proteins whose relative levels do not change is not a signature of the transition from specified to functional. It will be particularly interesting in the future to have occupancy information about phosphorylation during this period. It is reasonable that, because of the much lower mitotic index in these juvenile tadpole tissues and the decreased level of protein phosphorylation involved in gene expression, the absolute amount of phosphorylation in juvenile and adult tissues will be lower than in earlier embryos. However, these phosphorylations will most likely regulate important signaling and response circuits. Continual improvements in MS sensitivity should allow us to eventually investigate these potential regulatory circuits.

The data presented in this paper show that embryogenesis involves two different periods, the first being a period of generation of organization and differentiation, with minimal requirement for new protein. Only once this organized heterogeneity has been established do we see the second period, a period of biochemical differentiation where substantial change to the proteome does occur. The first period has been studied extensively by developmental biologists, attributing most of these changes to transcriptional regulation. The second period is ripe for attention.

## ACKNOWLEDGEMENTS

The authors enthusiastically thank R. Michael Gage for preparing the sample for mass spectrometry measurement, Rachael Jonas-Closs for excellent frog husbandry, and Alexander Lukyanov for writing the initial motif scoring code. This work was supported by NIH Grant OD award R24 OD031956 (to MWK and LP).

## CONTRIBUTIONS

Conceptualization (EV, MHW, LP, MWK), Experiments (LP, MW), Data curation (EV, MS, MK, LP), Formal Analysis (EV), Funding acquisition (LP, MWK), Investigation (EV, MHW, MS, MK, LP), Software (EV), Supervision (MHW, LP, MWK), Visualization (EV), Writing – original draft (EV), Writing – reviewing & editing (EV, MS, MK, LP, MWK).

## METHODS

### Illustrations

*Graphical abstract, Figure 1.A.* and *Figure 4.B.* illustrations © 2021 Natalya Zahn, <u>CC BY-NC 4.0</u> 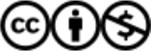.” The source of the illustrations is Xenbase (RRID:SCR_003280).

### Sample Preparation, Mass Spectrometry Measurement, and Preparing Data for Analysis

#### Samples

All handling of *Xenopus laevis* adults was carried out in accordance with HMS IACUC protocol IS00001365. *Xenopus laevis* J-line embryos were collected at Stages oocyte VI, egg, 9, 12, 18, 22, 24, 26, 30, 41, and 48. Embryos were de-jellied in 2% cysteine, pH 7.8, and flash frozen for later preparation. Stage VI oocytes were obtained by surgery from the same females from which eggs were collected for fertilization. Oocytes were harvested after the females had replenished their oocytes following the stimulation that resulted in collection of eggs.

#### Lysis

Frozen embryos/eggs at each stage were thawed and 5-6 uL of lysis buffer (1% NP-40, 250 mM sucrose (EMD cat# 8550), 10 mM EDTA (VWR cat# VW1474-01), 1 tablet Roche Complete mini protease inhibitor (cat# 11836153001) and 1 tablet Roche Phos STOP tablet (cat# 04906837001) per 10 mL, 25 mM HEPES (Sigma cat# H3375-500G) pH 7.2, 10 uM Combrestatin 4A (Santa Cruz cat# sc-204697), 10 uM Cytochalasin D (Santa Cruz cat# sc201442), 1 mM TCEP (ThermoFisher cat# 20490)) was added per embryo/egg on ice. The embryos/eggs were lysed by pipetting followed by thorough vortexing well. Preparation of the lysate from the Stage 48 tadpole required more extensive maceration of the sample with a pipette tip than did less mature samples.

#### Yolk removal

The yolk was removed from the samples by spinning at 4,000 xg for 4 minutes in a microcentrifuge. The lipids in the sample were then resuspended by lightly flicking the tube, being careful not to resuspend any of the yolk. The supernatant was transferred to another tube, and 2% SDS (Amresco cat# M112-500ML) was added. HEPES (Sigma cat# H3375-500G) (pH 7.2) was added to a final concentration of 100 mM. Additionally, during the yolk removal step material pelleted in addition to the expected yolk and pigment.

#### Proteomics sample prep*aration*

*The below proteomics sample preparation and mass spectrometry workflow was carried out for replicates A and B of protein and phosphorylation analysis (see Spreadsheet S4):*

#### Alkylation and cysteine protection

DTT (5 mM, pH ∼8.0, Sigma cat# 43819-1G) was added to each sample and the sample was incubated at 60 C for 20 minutes, then cooled to RT. Next, NEM (15 mM, Sigma cat# E3876-25G) was added to each sample and the samples were incubated for 20 minutes at room temperature. Finally, DTT (5 mM, pH∼8.0, Sigma 43819-1G) was added again and the samples were incubated at room temperature for 10 minutes.

#### Enzymatic Digestions

The proteins in the sample were precipitated using methanol/chloroform and were resuspended in 6 M Guanidine HCl (Sigma cat# G3272-1KG), buffered using 50 mM EPPS (Alfa Aesar cat# A13714), pH 9.0) to an estimated protein concentration of 5 mg/mL. The samples were heated to 60 C for 5 minutes and then allowed to cool to room temperature. The protein concentrations of each sample were determined by BCA assay (ThermoFisher cat# 23225). Next, each sample was diluted to 2 M Guanidine HCl with 5 mM EPPS (pH 9.0, Alfa Aesar cat# A13714), ensuring that the pH was at least 8.5. Lys-C (Wako Chemicals cat# 129-02541) was added at the higher of 1:100 w/w or 20 ng/uL and incubated for 12 hours at room temperature. The samples were diluted to 0.5 M Guanidine HCl with 5 mM EPPS (pH 9.0, Alfa Aesar cat# A13714), ensuring that the pH was at least 8.5. Lys-C (Wako Chemicals cat# 129-02541) was added again, as above, and allowed to digest for at least 15 minutes at room temperature. Additionally, Trypsin (Promega cat# V5113) was added at the higher of 1:50 w/w or 10 ng/uL and allowed to digest at 37 C for 8 hours. Samples were speed-vacuumed to dryness.

#### TMT labeling

Each sample (stage) was labeled using a distinct channel of TMT-11plex label (ThermoFisher cat# 90111 & A34807). Samples were resuspended in 500 mM EPPS (pH 8.0, Alfa Aesar cat# A13714), checking that the pH was close to 8.0, and incubated at 65 C for 5 minutes. After the samples returned to room temperature, fifteen uL of TMT reagent (20 mg/mL, ThermoFisher cat# 90111 & A34807) were added per each 100 ug of protein, and the reactions were incubated for 2 hours at room temperature. A small subset (∼1 ug/condition) was quenched and tested for TMT labeling efficiency, missed cleavage rate, and normalized total peptide count. Reactions were quenched by first heating the samples to 60 C for 5 minutes and then cooling to room temperature, followed by incubation with 0.5% hydroxylamine (Sigma cat# H9876) for 15 minutes at room temperature. The samples were then combined in another tube containing phosphoric acid (JT Baker cat# B34P0200) at 5% of the total combined volume. The combined sample was then speed-vacuumed to dryness.

#### Peptide purification, phosphopeptide enrichment and peptide fractionation

After TMT labeling, peptides were re-suspended in 1% formic acid and 0.1% trifluoroacetic acid (TFA) and purified by reversed phase chromatography using C18 Sep-Pak cartridges (Waters #WAT054945). After preconditioning the matrix with methanol and two full column washes with 1% formic acid with 0.1% TFA, peptides were bound by gravity flow. Columns were washed with 3 column volumes of 1% formic acid, peptides eluted with 95% acetonitrile in 1% formic acid and speed-vacuumed to dryness. Phosphopeptides were enriched by immobilized metal affinity chromatography (IMAC) using an Fe-NTA matrix (High-Select^TM^ Fe-NTA phosphopeptide enrichment kit, Thermo Scientific #A32992) according to manufacturer’s instructions. In brief, peptides were resuspended in loading buffer (0.1% TFA, 80% acetonitrile), allowed to bind to Fe-NTA matrix for 30 minutes, washed 2 times with loading buffer and once with water. Flow throughs of Fe-NTA columns were collected, pooled and peptides speed-vacuumed to dryness. Fe-NTA bound, enriched phosphopeptides were eluted with alkaline ammonia solution in water twice and dried down in parallel. Enrichment of phospho-peptides after TMT labelling avoids variability in the enrichment between time points and allows the protein level normalization scheme to be applied to the phospho-form data. The ability to use the same normalization scaling for each sample was especially important, since we expected large differences in bulk phosphorylation during our early time points due to the very high mitotic phosphorylation state of the egg [41,54,64].

Phospho-peptides were fractionated using alkaline reversed phase chromatography (Pierce^TM^ High pH Reversed Phase Peptide Fractionation Kit, Thermo Scientific #84868) into 12 fractions using increasing concentrations of acetonitrile from 10%-80% in basic binding buffer. Fractions 1 and 7, 2 and 8, 3 and 9, 4 and 10, 5 and 11, and 6 and 12 were combined. For fractionation of phospho de-enriched TMT labeled peptides, HPLC fractionation by alkaline reversed phase chromatography was carried out using an Agilent 1200 Series HPLC system with a flow rate of 600 ul/min over a 65-minute gradient with increasing concentrations of acetonitrile. 96 fractions of a 65-minute gradient of 13-44% buffer B (90% acetonitrile, 10mM ammonium bicarbonate, pH 8) in buffer A (5% acetonitrile, 10mM ammonium bicarbonate, pH 8) were collected and combined into 24 pooled fractions for MS analysis. Prior to injection, peptide samples were purified via Stage tips using punched out cookies of C18 3M^TM^ Empore^TM^ extraction disc material (Fisher Scientific #14-386-2) as matrix. Phospho-peptides were re-suspended in 1% formic acid prior to MS injection, non-phospho peptides in 1% formic acid with 5% acetonitrile.

#### Mass spectrometry measurement

Data were collected using a Multi Notch MS^3^ TMT method [17] using Orbitrap Fusion Lumos mass spectrometers (Thermo Fisher Scientific) coupled to a Proxeon EASY-nLC 1000 liquid chromatography (LC) system (Thermo Fisher Scientific). The 100 uM inner diameter nanospray capillary columns used were packed with C18 resin (2.6 μm, 150 Å, Thermo Fisher Scientific). Peptides of each fraction were separated over 4 and 5-hour acidic acetonitrile gradients for non-phospho peptides and 2.5 hours for phosphopeptides by LC prior to mass spectrometry (MS) injection. The scan sequence started with recording of an MS1 spectrum (Orbitrap analysis; resolution 120,000; mass range 400−1400 Th). MS2 analysis followed collision-induced dissociation (CID, CE=35) with a maximum ion injection time of 100-200 ms. For phospho-peptide measurement, a multi-stage activation method was used with a neutral loss of 97.9763/z. For TMT quantification, MS3 precursors were fragmented by high-energy collision induced dissociation (HCD) and analyzed in the Orbitrap at a resolution of 50,000 at 200 Th. Further details on the LC and MS parameters and settings used here were described previously [67].

#### Replicate C protein and phosphorylation sample preparation and mass spectrometry analysis was carried out as above for A and B with the following differences

Peptides were alkylated with iodoacetamide instead of N-ethylmaleimide (NEM) and labeled with TMT-Pro 16-plex reagents (Thermo Fisher, Cat# A44520). Mass spectrometry measurements were acquired on an Orbitrap Eclipse mass spectrometer (Thermo Fisher Scientific). For protein measurements, the mass spectrometer was equipped with a FAIMS-Pro interface, and three different CVs (−51, −67, −83) were analyzed sequentially in the same run for all MS1 and MS2 scans [68]. The scan sequence began with acquisition of an MS1 spectrum (Orbitrap analysis; resolution 120,000; mass range 400–1100 Th; 5e5 minimum intensity threshold; 60-second precursor exclusion). Precursors were then fragmented and analyzed with an MS2 spectrum (Orbitrap analysis; resolution 50,000; 0.7 Th isolation window; 37 normalized HCD in fixed collision mode; first mass of 110 Da; maximum injection time of 86 ms). Phosphorylation measurements were performed with a single CV for each phosphorylation sample. Individual CV runs were done using −32, −42, −52, −62, −72, or −82. An additional run employed the same triple CV method as the protein analysis (−51, −67, −83).

#### Mapping and normalization

For mapping of protein and phospho data (peptide-spectra matches for MS), we used, as a main database of reference, sequences from the *X. laevis* genome assembly (v9p2 of assembly: a total of 46,582 sequences) downloaded from Xenbase [69] (https://download.xenbase.org/xenbase/Genomics/JGI/ Xenla9.2/sequences/XENLA_9.2_Xenbase.pep.fa.gz) combined with 13 mitochondrial protein sequences. Peptides were searched with a COMET-based in-house software with a target decoy database strategy and a false discovery rate (FDR) of 2% set for peptide-spectrum matches by linear discriminant analysis (LDA). The peptide and phospho-peptide identifications from both replicates were then combined for a final collapsed protein-level FDR of 2%. Phospho-peptide localization confidence was quantified using the AScore method. All analyzed phospho-forms have AScore values >=13. For Replicates A and B, quantitative information on peptides and phospho-peptides was derived from MS3 scans. Details of the TMT intensity quantification method and further search parameters applied were described recently [67]. For Replicates C, quantitative information on peptides and phospho-peptides was derived from MS2 scans (see above). Only peptides with a sum of TMT s/n >= 200 were used. The channels were further normalized assuming that the same amount of protein per time point was combined after multiplexing, using total time point signal as a proxy for protein amount.

Phospho-forms are distinct peptides with distinct phosphorylation events that could have a single phosphorylation or multiple phosphorylations. More than 80% of measured phospho-forms in this dataset have only one phosphorylation (*Figure S1.A.*). We identified all measured peptides that had the same residues phosphorylated. In the cases where multiple versions exist (because of internal lysines and arginines), we used only the most fully cleaved version of the peptide.

#### Assigning Human Gene Symbols to Xenopus laevis references

We used the mappings between the X. laevis 9.2 reference gene models to human gene symbols reported in [70]. Due to a partial genome duplication in the pseudo-tetraploid *Xenopus laevis*, many proteins have homoeologous genes (sometimes termed alloalleles), which have separate gene references [71]. These alloalleles are analyzed separately.

### Data Accessibility

Mass spectrometry spectra files have been deposited to the ProteomeXchange Consortium via the PRIDE [72] partner repository with the dataset identifier PXD060481. The set of raw files associated with each replicate can be found in *Spreadsheet S4*. The mapped and normalized relative protein, absolute protein, and relative phospho datasets are included as Spreadsheets 1, 2, and 3, respectively. Python code for analyzing the spectra search output can be found here: https://github.com/elizabeth-van-itallie/DevSeries_Processing.

### Over Representation Analysis (ORA)

#### Relative Protein

We employed a two-step approach. First we used the WebGESTALT web server (www.webgestalt.org) [45,73] to perform ORA with the “Biological Process nonRedundant’’ database. For each cluster, we used the assigned human gene symbols for the proteins in that cluster as the “gene list” and used all of the human gene symbols assigned to proteins quantified as the “background”. The set of human gene symbols for the measured proteins is not unique, both because of allo-alleales and cases where multiple Xenopus proteins map to the same human protein, and not all replicate gene symbols are in the same “gene lists’’ that are tested for overrepresentation. The complete set of overrepresented GO gene sets for all clusters with FDR <= 5% is reported in *Spreadsheet S1*. For the subset of gene sets shown in Figure 2B, we identified all of the clusters for which the gene set was over-represented. Then we determined the p-value of the hypergeometric test for each cluster in which the gene set is over-represented. If the p-value is less than or equal to 0.05, it is visualized with a circle of area proportional to the -log10 of the p-value.

#### Relative Phospho (Phospho-proteins)

Similar to above, for each cluster, we used the assigned human gene symbols for the phospho-proteins in that cluster as the “gene list” and used all of the phospho-protein human gene symbols as the “background.” A small number of the GO gene sets that’s have FDR <= 5% are shown in Figure 5C, and the complete set of overrepresented GO gene sets for all clusters with FDR <= 5% is reported in *Spreadsheet S2*.

#### Relative Phospho (Kinase/Phospho-site)

For clusters B and D, where there were both over-represented motifs and a substantial number of phospho-sites, we used the set of phospho-sites that have human phospho-site matches as the input PTM list (see *Identifying homologous phospho-acceptors between Xenopus and human using global and local sequence homology*), and we used the set of all phospho-sites that have human phospho-site matches as the reference set. The top-10 over-represented sets for both clusters are reported in *Spreadsheet S3*.

### Gene Set Enrichment Analysis (GSEA)

We used the WebGESTALT web server (www.webgestalt.org) [45,73] to perform GSEA for the fraction of maximum fold change before the tadpole metric. The metric was calculated for each protein quantified in the dataset and assigned human gene symbols were used. The full list of positively and negatively enriched gene sets is shown in *Figure 3.C*.

### Measuring average total protein amount per embryo

Sets of ten embryos from the same clutch were flash frozen at the same developmental stages as the proteomics dataset, except that Stage VI oocytes were not collected. The egg through Stage 41 samples were prepared exactly as described above under **Sample Preparation and Mass Spectrometry Measurement**. An additional egg sample and the Stage 48 sample were lysed in the same lysis buffer but with SDS already added to 2%, and there was no yolk spin out. After cysteine protection, and before the methanol chloroform precipitation, the concentration of a 1:10 dilution of the lysate (1:50 for the egg with yolk) was measured with a reducing agent compatible BCA assay (ThermoFisher 23252).

### Estimating Absolute Protein Amounts for Individual Proteins

Using an updated mapping of the deep egg proteome absolute abundance dataset [2,70], proteins that are measured in both datasets were used to estimate the linear relationship between the log of the fraction of the MS1 signal attributed to the egg and the log of estimated protein concentration in the egg from the PHROG database reference [2]. The Python3 “scipy.odr” package was used to perform the orthogonal regression because there is error in both dimensions. The resulting fit and the assumption that the volume of non-yolk cytoplasm in the egg is 0.33 uL were used to estimate the number of moles of protein in the egg for all measured proteins. These values were then scaled by the relative protein change data and the relative change in absolute amount of total protein in the embryo, to determine the number of moles at the other stages. It was assumed that the total amount of protein in the oocyte and egg is the same.

### Protein Localization Classification

Localization annotations for nucleus, membrane, secreted, and cytoplasm were downloaded from the manually annotated and reviewed SwissProt database via UniProt. The assignments for all measured proteins are reported in the “Localization” column of Data set 2.

### Tissue Specificity

Proteins were classified as tissue-specific if they were both (1) not measured in the egg proteome database [2], and (2) if the matched human gene symbol was classified as “Tissue enriched” or “Group Enriched” in the Tissue Atlas Database [38]. Whether a protein was measured in the egg proteome reference is reported in the “InDeepEgg” column of Data set 2, and the Tissue Specificity classification is reported in the “TissueSpecific” column of Data set 2.

### Identifying homologous phospho-acceptors between Xenopus and human using global and local sequence homology

First we identified the human sequence in the reference set of human protein sequences that is the best match for each of the Xenopus protein references on which we have measured a phospho-site. We used the human-phosphosite-fastas.fasta from Phosphosite.org after filtering to remove all isoform references [16]. We chose the BLASTP result with the smallest e-value, and only used alignments if the e-value is less than 1e^-20^. For all the phosphorylated residues on each Xenopus reference, we used the alignment to determine if there is a corresponding potentially phosphorylated residue that aligns on the matched human reference. At this step there are five possible outcomes from evaluating the alignments at each queried residue position: (1) aligned to an identical human residue *(match)*; (2) S->T or T->S mismatch *(match)*; (3) aligned to a different human residue *(mismatch)*; (4) aligned to a gap in the human reference *(mismatch)*; and (5) found on a part of the *Xenopus laevis* reference that does not align to the human reference *(mismatch)*. We considered serine and threonine residues that align to the opposite residue as matches, for two reasons: (1) mutations between S and T are relatively common over evolutionary time; and (2) there are many kinases that phosphorylate both serine and threonines. In the cases where the Xenopus residue aligns to a human residue, we determine the six amino acids that flank the aligned residue N- and C-terminally (the “motif”). Then we score the local sequence homology for all these aligned motifs. The motif score is based on the Blosum 90 substitution penalty matrix. The alignment score is calculated from the flanking amino-acids only, not the aligned amino-acids. Since the alignment score depends not only on the number of matches, but also on the identities of the constituent amino acids, we normalize for sequence effects of the score by finding the “best score” (largest of human or xenopus alignment score to itself) and “worst score” (smallest of human or xenopus alignment score to its flipped self). The “motif score” is the Xenopus against human alignment score minus the “worst score” divided by the “best score” minus the worst score. ( *Motif Score = ( Xenopus: Human alignment - “worst score”) / ( “best score” - “worst score”)*). Then, using the alignments to non-phosphorylatable residues as our “false discoveries,” we determined the false discovery rate as a function of motif score. We retain matches that have motif scores greater than or equal to 0.7, which means that we have an FDR of ∼10%. For the python3 based snakemake pipeline used for this analysis see: https://github.com/elizabeth-van-itallie/phospho_matching.

### Motif Enrichment

Motif enrichment was performed using the R motif-x implementation [42,43]. For each of the eight k-means clusters, the +/- 6 amino acid motifs for all phospho-forms in the clusters were the foreground set and the +/- 6 amino acid motifs for all of the phospho-forms in the dataset were used as the background. The enrichment was performed with the central residue as Serine and Threonine for all clusters; thus, sixteen total tests were performed. The min.seqs parameter was set to 20 and the pval.cutoff parameter was set to 1e^-6^. Where multiply phosphorylated peptides had multiple motifs per phospho-form, we considered each motif separately.

