## Supplementatry materials for "Transitions in the proteome and phospho-proteome during *Xenopus laevis* development"

SPREADSHEET COLUMN DESCRIPTIONS and SUPPLEMENTAL FIGURE LEGENDS and SUPPLEMENTAL TABLE and SUPPLEMENTAL FIGURES for:

**Main Spreadsheets**

**Spreadsheet 1:** Relative Protein Trends

The columns are as follows:
**Index column:** The gene name from the Xenopus laevis protein reference(V9.2).
**laevisGene:** The Xenopus laevis gene name.
**humanGene:** The matched human gene name.
**description:** Brief description of the humanGene.
**Chan_[1-11]_A:** Normalized relative protein signal data if the protein was measured in Replicate A. The Channel numbers correspond in order to the collected timepoints shown in Figure 1.A.
**Chan_[1-11]_B:** Same as above but for Replicate B.
**Chan_[1-11]_C:** Same as above but for Replicate C.
**Chan_[1-11]_Median:** The median of the Replicate trends of all the replicates where that protein was measured.
**Yes[A,B,C]:** Logical index for whether the protein was measured in that replicates (1 = measured, 0 = not measured)
**pccAB:** Pearson Correlation Coefficient for Replicate A and Replicate B.
**pccAC:** Pearson Correlation Coefficient for Replicate A and Replicate C.
**pccBC:** Pearson Correlation Coefficient for Replicate B and Replicate C.
**pcc_Median:** The median of the Pearson Correlation Coefficients of the Pearson Correlation Coefficients for the pairs of measured data.
**Cluster:** The cluster (shown in Figure 2A) that the protein median trend is in.
**ffcA:** The fractional change metric (introduced in Figure 3) for the Replicate A data.
**ffcB:** Same as above but for Replicate B data.
**ffcC:** Same as above but for Replicate C data.
**ffcMedianTrend:** Same as above for the median trend.
**ProteinClass:** Either TF (transcription factor), E3 (E3 ubiquitin ligases), Kinase, Receptor, Ligand or not assigned (See Figure 3B).

**Spreadsheet 2:** Absolute Protein Amounts

The columns are as follows:
**Index column (Reference_v9p2):** The gene name from the Xenopus laevis protein reference (V92).
**laevisGene:** The Xenopus laevis gene name.
**humanGene:** The matched human gene name.
**description:** Brief description of the humanGene.
**Chan_[1-11]_A:** Estimated protein amount in picomoles if the data was measured in Replicate A. Then channel numbers correspond in order to the collected timepoints shown in Figure 1.A.
**Chan_[1-11]_B:** Same as above but for Replicate B.
**Chan_[1-11]_C:** Same as above but for Replicate C.
**Chan_[1-11]_Median:** The median of the Replicate trends of all the replicates where that protein was measured.
**Localization:** Cell localization categories introduced in Figure 4. Options are “NoData,” “Nuclear,” “Cytoplasm,” “Membrane,” and “Secreted.”
**Diff_30to41_A:** The difference between Channel_10_A and Channel_9_A, where a positive value means that Chanel_10_A is larger than Channel_9_A, and a negative value is the opposite.
**Diff_30to41_B:** Same as above but for Replicate B.
**Diff_30to41_C:** Same as above but for Replicate C.
**Diff_41to48_A:** The difference between Channel_11_A and Channel_10_A, where a positive value mea ns that Chanel_11_A is larger than Channel_10_A, and a negative value is the opposite.
**Diff_41to48_B:** Same as above but for Replicate B.
**Diff_41to48_C:** Same as above but for Replicate C.
**InDeepEgg:** Whether the protein was measured in Wühr et al. 2014 (1 means YES, 0 means NO).

**TissueSpecific:** Classification of tissue specificity used for Figure 4.E. Options are “NoData,” “Yes,” and “No.”

**Spreadsheet 3:** Relative Phosphorylation Trends

The columns are as follows:

**PhosphoFormIndex:** Index column

**Reference_v9p2:** The gene name from the Xenopus laevis protein reference (V92).

**laevisGene:** The Xenopus laevis gene name.

**humanGene:** The matched human gene name.

**description:** Brief description of the humanGene.

**Site_Positions:** The residue number(s) on the reference that have been determined to be phosphorylated. If there is more than one measured phosphorylation, the numbers are separated with semi-colons.

**Summary_Start_Pos:** The most N-terminal residue number for the spanning set of peptides associated with the phosphorylation site(s).

**Summary_End_Pos:** The most C-terminal residue number for the spanning set of peptides associated with the phosphorylation site(s).

**Motifs_Clean:** +/- 6 motifs around the residues in “Site Positions.” If there is more than one site than the motifs are semi-colon delimited.

**Data_A:** Flag for if the phospho-form was measured in replicate A (1 if yes, 0 if not).

**Data_B:** Same as above but for Replicate B.

**Data_C:** Same as above but for Replicate C.

**Human_Reference:** The human best match reference that was found for the phospho-site mapping described in Figure 5 and Supplemental Figure 15.

**Human_Residue:** The human matched residue on the reference in Human_Reference if there is a match that passes the matching criteria.

**Human_Motif:** The +/- 6 motifs from the Human reference sequence for the matched Human residues.

**For each replicate A, B, C there are the following columns:**

**Peptide:** The sequence of the measured phospho-form peptide where the @ signal designates the phosphorylation.

**Site_Scores:** The score associated with the localization of the phosphorylation where very large scores are highly confident (e.g. there is only one phosphorylatable residue in the peptide) and lower scores reflect lower confidence.

**Start_Position:** The residue number of the N-terminal end of the measured peptide.

**End_Position:** The residue number of the C-terminal end of the measured peptide.

**Chan_[1-11]:** The relative phospho-form signal normalized normalized across the channels such that the median is 1. There is no normalization to the protein trend on which this phospho-form has been measured.

**Chan_[1-11]_Median:** The median phospho-form relative signal values for each channel across the measured replicates.

**Chan_[1-11]_Median_Prot:** The median relative signal values for the protein reference that the phospho-form is measured on. These values are the same as the Chan_[1-11]_Median values in Data set 1.

**CoCluster:** The co-cluster in Figure 5.B. that the median phospho-from and median protein trends are assigned to.

**PPCorr_A:** The PPCorr value (introduced in Supplemental Figure 13.A.) for Replicate A. This value uses the phospho-form signal measured in Replicate_A and the relative protein data for Replicate_A. If there is no data for either of these measurements, then the value for this column is NaN.

**PPCorr_B:** Same as above but for Replicate B.

**PPCorr_C:** Same as above but for Replicate C.

**PPCorr_Median:** The median of the three values above. PPCorr is not calculated using the median phospho-form and protein trends.

**Phos_FC_A:** The Phos_FC value (introduce in Supplemental Figure 13.B.) for Replicate A.

**Phos_FC_B:** The same as above but for Replicate B.

**Phos_FC_C:** The same as above but for Replicate C.

**Phos_FC_Median:** The median of the three values above.

**Motif_Class:** For single sites, the motif class (as defined in Figure 6.A) for the motif.

**Phospho-Acceptor:** For single sites, the phospho-acceptor residue (S,T, or Y).

**PCC_AB:** The Pearson Correlation Coefficient of the phospho-form signals of Replicates A and B.

**PCC_AC:** The Pearson Correlation Coefficient of the phospho-form signals of Replicates A and C.
**PCC_BC:** Pearson Correlation Coefficient of the phospho-form signals of Replicates B and C.

#### Supplemental Figures

**Figure S1:** Properties of the measured phospho-proteome.

**A.** Fraction of phospho-forms measured with one, two, or three phosphorylations per peptide.

**B.** Cumulative fraction of phospho-forms measured per phospho-protein. Approximately 50% of phospho-proteins have one phospho-form.

**Figure S2:** Proteomic and phospho-proteomics measurement reproducibility.

**A,B.** Pairwise Pearson correlation coefficients between replicates at each developmental stage for both protein data (A) and phosphorylation data (B). Comparing replicates A and B, for all stages, the same stage from the other replicate has the largest Pearson correlation coefficient. Comparing replicates A and B with replicate C, in both cases the Stage 26 timepoint in replicate C is closest to the Stage 24 timepoint in replicates A and B. The Stage 30 timepoint in replicate C is also closest to the Stage 26 timepoint in replicate A rather than to the Stage 30 timepoint in replicate A. Thus, replicates A and B are more similar to each other than they are to replicate C, and part of the replicate C timeseries is shifted earlier relative to replicates A and B.

**C.** Distribution of Pearson correlation coefficients for all replicate pairwise comparisons of the protein (left) and phosphorylation (right) time series compared to a background of 10,000 scrambled matches.

**Figure S3:** The original twelve k-means relative protein clusters before merging.

The median trend (black line) and ten to ninety percentile regions (gray shading) are shown for the twelve-original k-means relative trend protein clusters. The mapping from the merged clusters to the original clusters is as follows: 1:1, 2:2, 3:3, 4:4, 5:{5,6}, 6:{7,8}, 7:{9, 10, 11}, 8:12.

**Figure S4:** Individual replicate relative protein signal corresponding to the median trends shown in Figure 2C. If a trend is not shown for a replicate, then it was not measured in that replicate.

**A.** Replicate trends for the proteins shown in Figure 2.C.i.

**B.** Replicate trends for the proteins shown in Figure 2.C.ii.

**C.** Replicate trends for the proteins shown in Figure 2.C.iii.

**Figure S5:** Schematic examples of the Initial Development Maximum Fold Change and the Across Development Maximum Fold Change. The Initial Development Maximum Fold Change (red) is the maximum fold change between any two timepoints between and including the oocyte and Stage 30. The Across Development Maximum Fold Change (grey) is the maximum fold change between any two timepoints in the entire time series. Examples are shown where the Initial Dev. Max F.C. is less than the Across Dev. Max. F.C. (~0.4) **(left)**, and where the Initial Dev. Max. F.C. and the Across Dev. Max. F.C. are equal (~1.0) **(right)**.

**Figure S6:** Violin plots of the replicate trend FFCs for the six different classes and all proteins (center black line = median; top and bottom black lines = max and min, respectively). The numbers below the violin plots are the number of proteins in the classes for that replicate. For replicate A, medians of transcription factors (TFs), E3 ubiquitin ligases (E3s), kinases, and ligands are statistically significantly different than that of all proteins ($p < 0.001$; Kruskal Wallis test, Bonferroni critical value for multiple comparisons - all comparisons in Figure 3.B and in this supplemental figure - with post-hoc Dunn’s test). For replicate B, medians of transcription factors (TFs), E3 ubiquitin ligases (E3s), and ligands are statistically significantly different than that of all proteins (same statistical methods as A). For replicate C, medians of transcription factors (TFs), E3 ubiquitin ligases (E3s), kinases, and ligands are statistically significantly different than that of all proteins (same statistical methods as B).

**Figure S7:** Individual replicate relative protein signal corresponding to the median trends shown in Figure 3Bii. If a trend is not shown for a replicate, then it was not measured in that replicate.

**Figure S8:** Individual replicate relative protein signal corresponding to the median trends shown in Figure 3Cii. If a trend is not shown for a replicate, then it was not measured in that replicate.

**Figure S9:** 86.5% of the protein in the egg is yolk; only 13.5% of the protein in the egg is not yolk. The means of three technical replicates are shown with standard error of the mean.

**Figure S10:** Data and linear fit for the relationship between ion current and protein concentration in the egg for the three replicates.

**Figure S11:** Individual replicate absolute protein amount trends corresponding to the median trends shown in Figure 4D (A) and Figure 4F (B).

**Figure S12:** Examples of changes in the absolute abundance of tissue specific cytoplasmic proteins that are expressed in muscle tissue.

**Figure S13:** Schematics illustrating two metrics for comparing the properties of the phospho-form protein co-clusters using the examples introduced in Figure 5.A.

**A.** PP Corr is the Pearson Correlation Coefficient of the phospho-form trend and the trend of the protein that the phosphorylation is on. The phospho-form and protein example Figure 5.A.iii. has a low PP Corr value because the protein and phospho-form trends are different and the magnitude of change for the phospho-form change is large. In Figure 5.A.ii., the PP Corr value is moderate because the trends are similar but the magnitude of change for the trends is small. In Figure 5.A.i., the PP Corr value is high because the phospho-form and protein trends are very similar and there is substantial magnitude change.

**B.** Phos FC is the maximum fold change between any two timepoints in the phospho-form timeseries. The phospho-form and protein example Figure 5.A.ii. has a low Phos FC. For Figure 5.A.i. the Phos FC value is moderate, and for Figure 5.A.iii. the Phos FC value is high.

**Figure S14:** The original twelve relative signal protein and phospho-form k-means co-clusters before merging (black and gray is protein, purple and pink is phospho-form). The median trend (black line and purple line) and ten to ninety percentile regions (gray and pink shading) are shown for the twelve-original k-means relative trend co-clusters. The mapping from the merged co-clusters to the original clusters is as follows: 1:{4,11}, 2:6, 3:8, 4:{2,3,9}, 5:1, 6:7, 7:12, 8:{5,10}.

**Figure S15:** A custom pipeline to identify homologous phospho-residues across species to take advantage of human-centric databases and tools. The snakemake code for this pipeline can be found here: https://github.com/elizabeth-van-itallie/phospho_matching.

**A.** Overview of the steps of the pipeline as outlined in Methods.

(left) Identifying the homologous human protein with BLASTP and e-value filtering.

(center) The alignment of the homologous proteins is assessed to determine what the phosphorylated Xenopus residues align to in the human sequence.

(right) Similarity of the local sequence (+/- 6 aa) is quantified with a BLOSSUM-90 based motif score to differentially weight mismatches.

**B.** Histogram of the alignment results for the 12,442 phosphorylated Xenopus residues. ”Match” means that the Xenopus phosphorylated residue aligns to the same residue in the best match human sequence, “S/T Swap” means that the Xenopus phosphorylated residue aligns to a different reside, but one that can phosphorylated, “Not Phosphorylatable” means that the Xenopus phosphorylated residue aligns to a residue that cannot be phosphorylated, “Gap” means that the Xenopus phosphorylated residue aligns to a gap in the human best match protein, “No Residue” means that the Xenopus phosphorylated residue did not align at all to the human sequences, and “No Reference” means that there is no human sequence that meets the criteria.

**C.** To determine the appropriate threshold for our motif score we compared the distribution of motif scores for matched residues (including S/T swaps) to motif scores for the aligned Xenopus/human mismatches (human residue is not a phospho-acceptor, “Not Phosphorylatable”). We chose an FDR of 10% which resulted in a motif score cutoff of >= 0.7. Residues with motif scores above this threshold are considered to have homologous human residues where the combination of global and local sequence homology allows us to carry over information about these human residues to Xenopus.

**Figure S16: Putative CK2 substrates in Co-cluster B.**

**A.** Two putative CK2 substrates in co-cluster B (Figure 5). The solid lines are the median protein trends and then dashed lines are the phospho-form trends. The Xenopus motif and the phospho-residue and motif for the best matched human sites are also shown.

**B.** Individual replicate data for the HSP90AB1 phospho-form shown in A (left).
**C.** Individual replicate data for the PSMA3 phospho-form shown in A (right).

**Figure S17: Individual replicate phospho-form and protein data for the median trends in Figure 5.E.**

**A.** CDK1 protein (solid) and double phosphorylation T-45 and Y-46 (dashed) was measured in replicates A and C.

**B.** YBX1 protein (solid) and phosphorylation on S-153 (dashed) was measured in replicates A, B, and C.

**C.** PRKAR1A protein (solid) and phosphorylation on S-82 (dashed) was measured in replicates A, B, and C.

**Figure S18: Underlying per replicate data for the summary data shown in Figure 6.C.** Box plots without outliers are shown for all the categories of all measured phospho-forms in the replicate: all single phosphorylation phospho-forms, proline directed motifs, acidic motifs, basic motifs, all sites that do not have proline-directed, acidic or basic motifs (“other”), serine phospho-forms, and threonine phospho-forms. The numbers under the labels are the number of single phospho-forms in that class for that replicate.

**Figure S19: Comparison of the distributions of Median PhosFC values for Serine and Threonine single phosphorylated residue phospho-forms.**

**Figure S20: Individual replicate phospho-form and protein data for the median trends in Figure 6.D.**

**A.** Replicate trends for the two phospho-forms and protein trends in the left main panel (BMS1 top, MRTO4 bottom).

**B.** Replicate trends for the two phospho-forms and protein trends in the center main panel (STK24 left, STK26 right).

**C.** Replicate trends for the two phospho-forms and protein trends in the right main panel (MOB1B left, STK3 right).

**Supplemental Table 1.** Complete results for the testing for enrichment of motifs in the phospho-from co-clusters shown in Figure 5.D.

| Cluster | Central Residue | Enriched Motifs (p <= 1e-6, ordered by counts in the cluster) |
|---|---|---|
| A | S | None |
| A | T | None |
| B | S | ......SD.....<br>......SE.....<br>......SS..... |
| B | T | None |
| C | S | ......S..E...<br>..G...S...... |
| C | T | None |
| D | S | ......SP.....<br>......S..K...<br>...L..S......<br>....F.SP..... |
| D | T | ......TP..... |
| E | S | None |
| E | T | None |
| F | S | None |
| F | T | None |
| G | S | None |
| G | T | None |
| H | S | None |
| H | T | None |

**Figure S1**

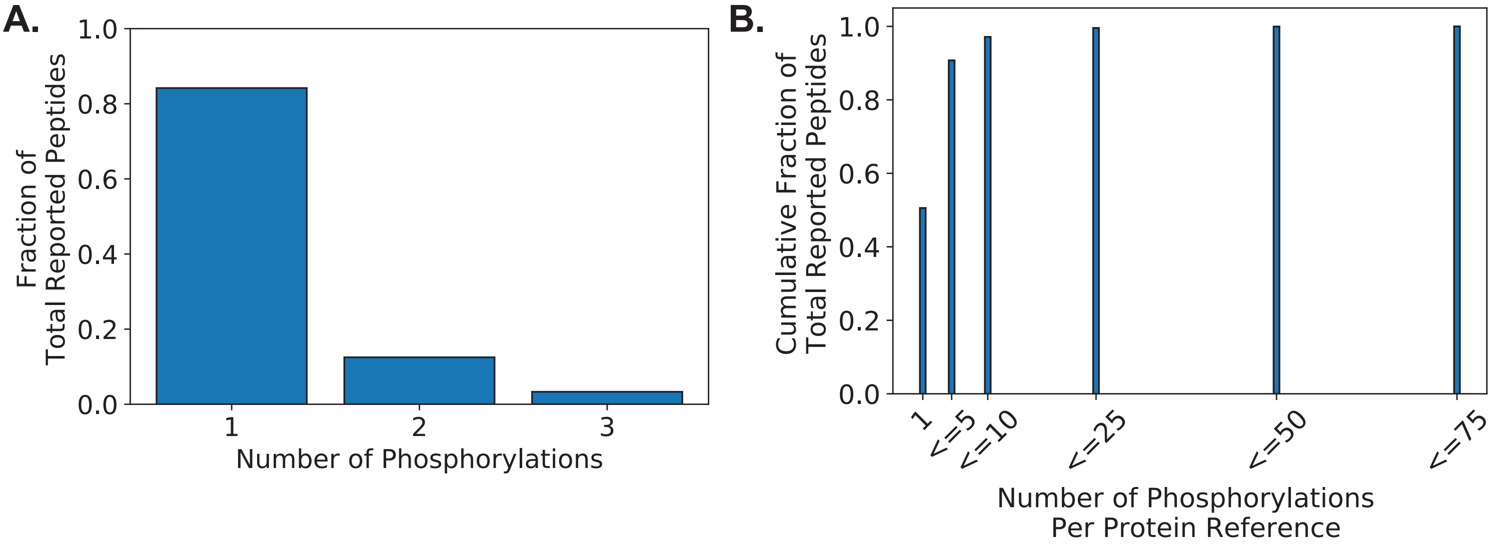
A.
1.0
0.8
0.6
0.4
0.2
0.0
Fraction of
Total Reported Peptides
1
2
3
Number of Phosphorylations
B.
1.0
0.8
0.6
0.4
0.2
0.0
Cumulative Fraction of
Total Reported Peptides
1
<=5
<=10
<=25
<=50
<=75
Number of Phosphorylations
Per Protein Reference

### Figure S2

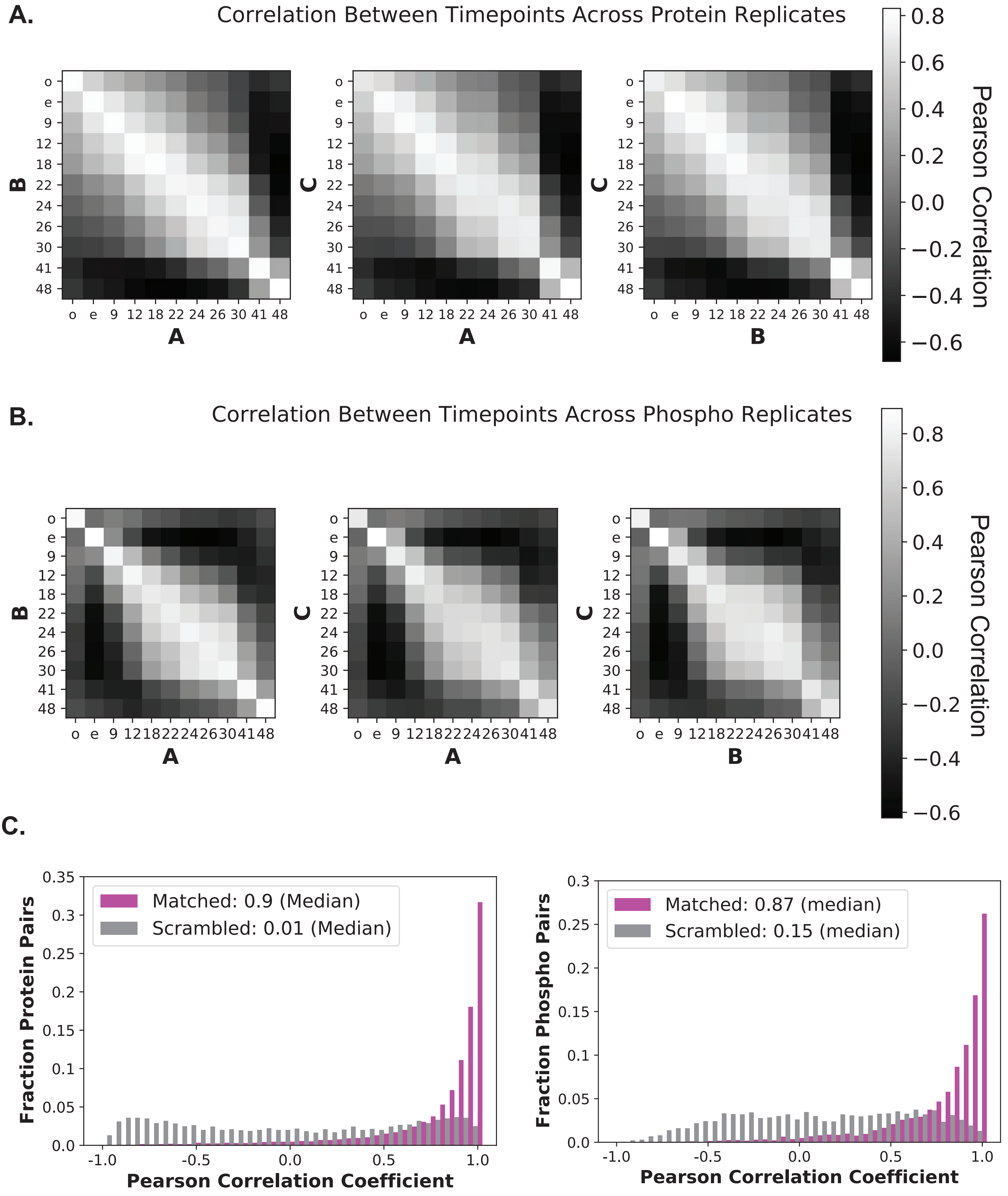
A.
Correlation Between Timepoints Across Protein Replicates
Pearson Correlation
B.
Correlation Between Timepoints Across Phospho Replicates
Pearson Correlation
C.
Matched: 0.9 (Median)
Scrambled: 0.01 (Median)
Fraction Protein Pairs
Pearson Correlation Coefficient
Matched: 0.87 (median)
Scrambled: 0.15 (median)
Fraction Phospho Pairs
Pearson Correlation Coefficient

### Figure S3

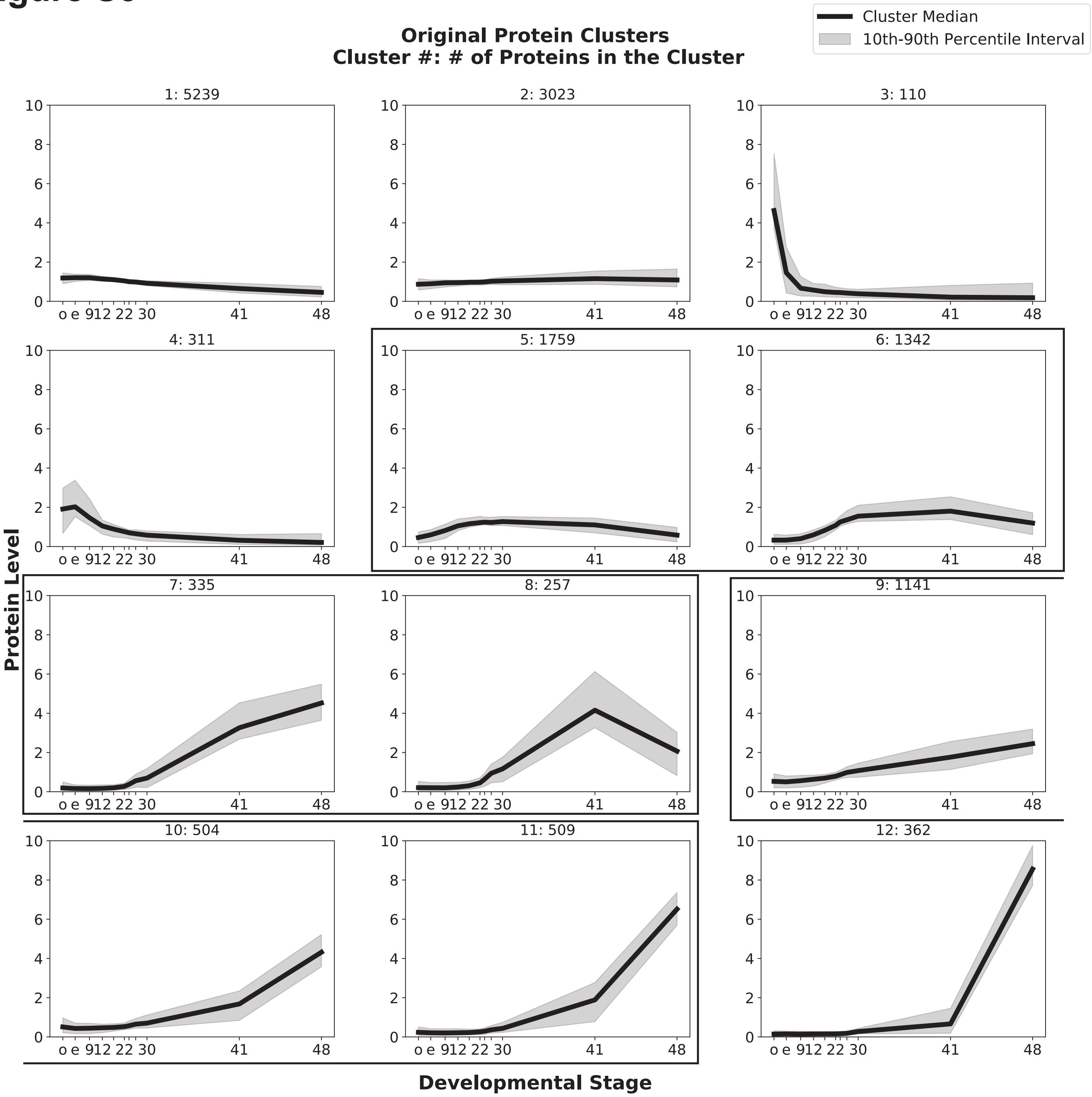
Cluster Median
10th-90th Percentile Interval
Original Protein Clusters
Cluster #: # of Proteins in the Cluster
1: 5239
2: 3023
3: 110
4: 311
5: 1759
6: 1342
7: 335
8: 257
9: 1141
10: 504
11: 509
12: 362
Protein Level
Developmental Stage
o e 912 22 30 41 48

### Figure S4

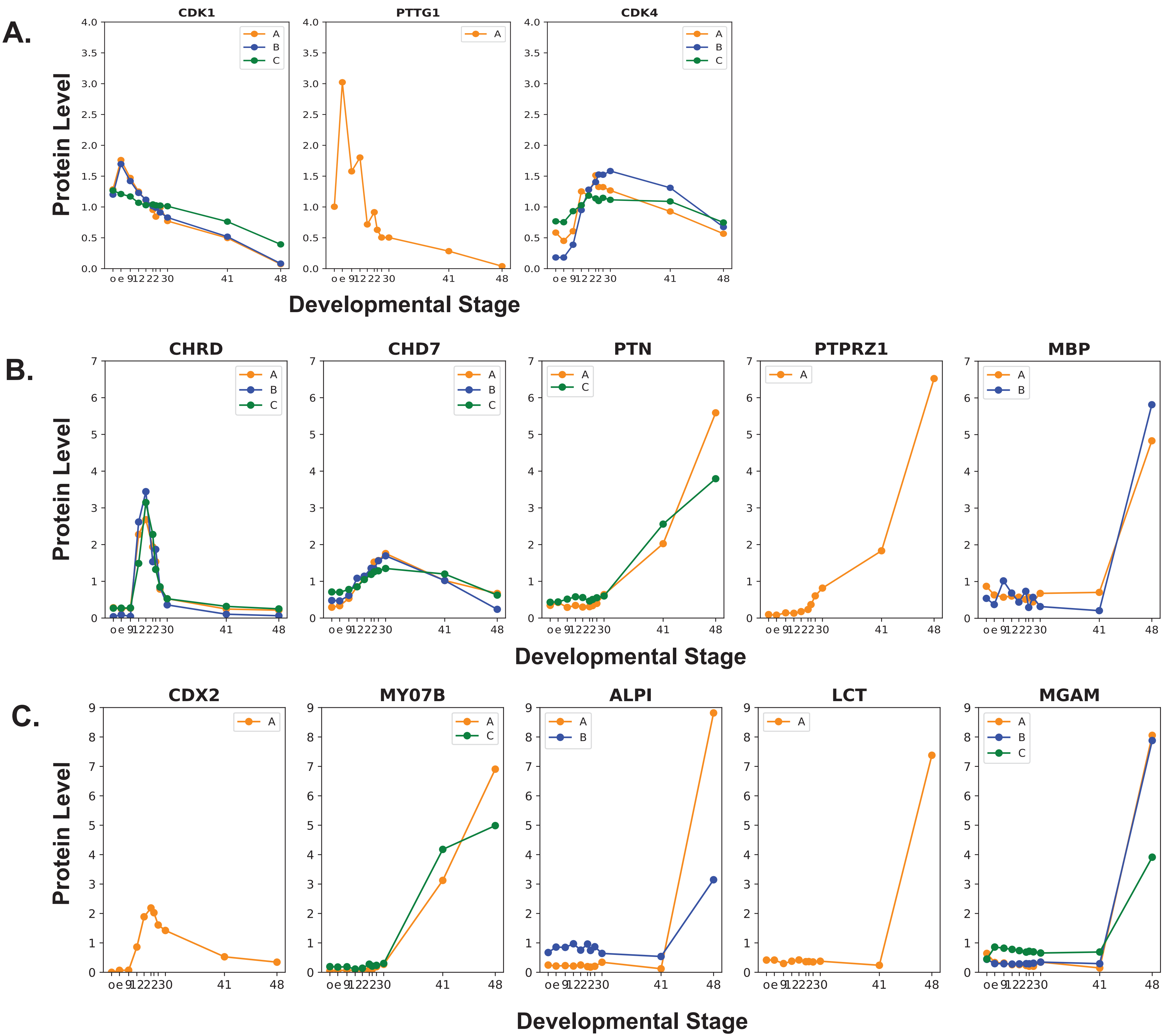
A.
B.
C.
CDK1
PTTG1
CDK4
CHRD
CHD7
PTN
PTPRZ1
MBP
CDX2
MY07B
ALPI
LCT
MGAM
Protein Level
Developmental Stage
A
B
C

### Figure S5

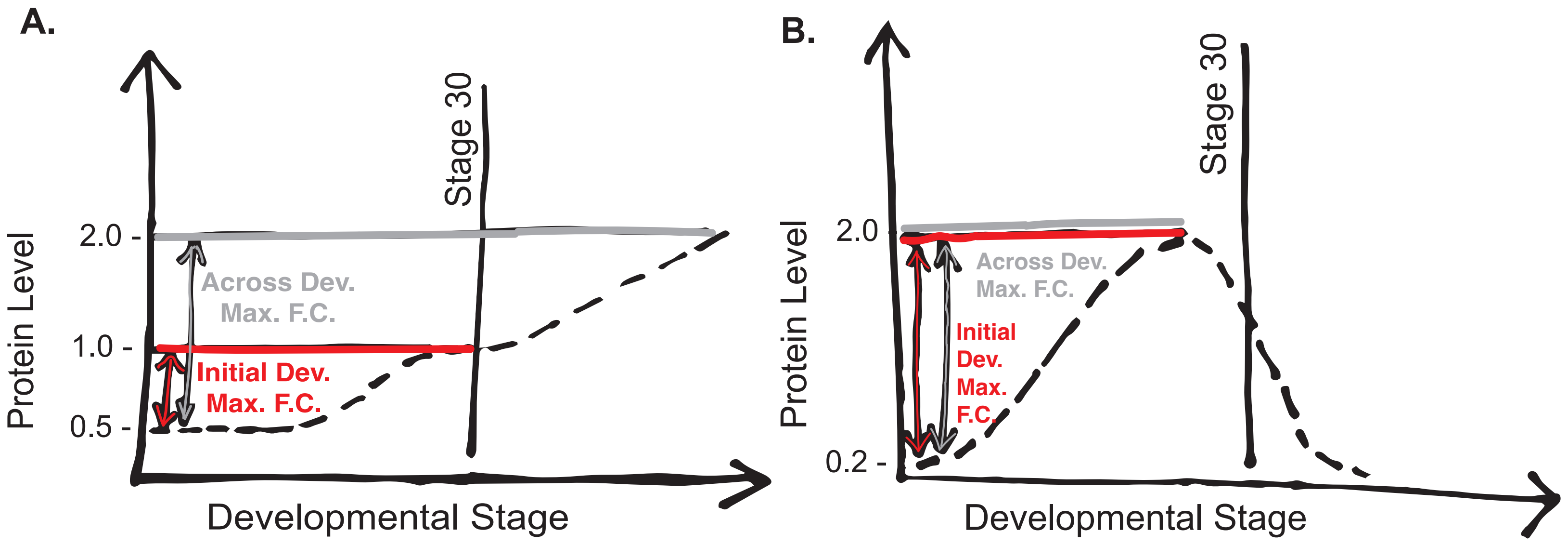


FFC = Initial Dev Max FC / Across Dev Max FC
= 2 / 4
= 0.5

FFC = Initial Dev Max FC / Across Dev Max FC
= 10 / 10
= 1

### Figure S6

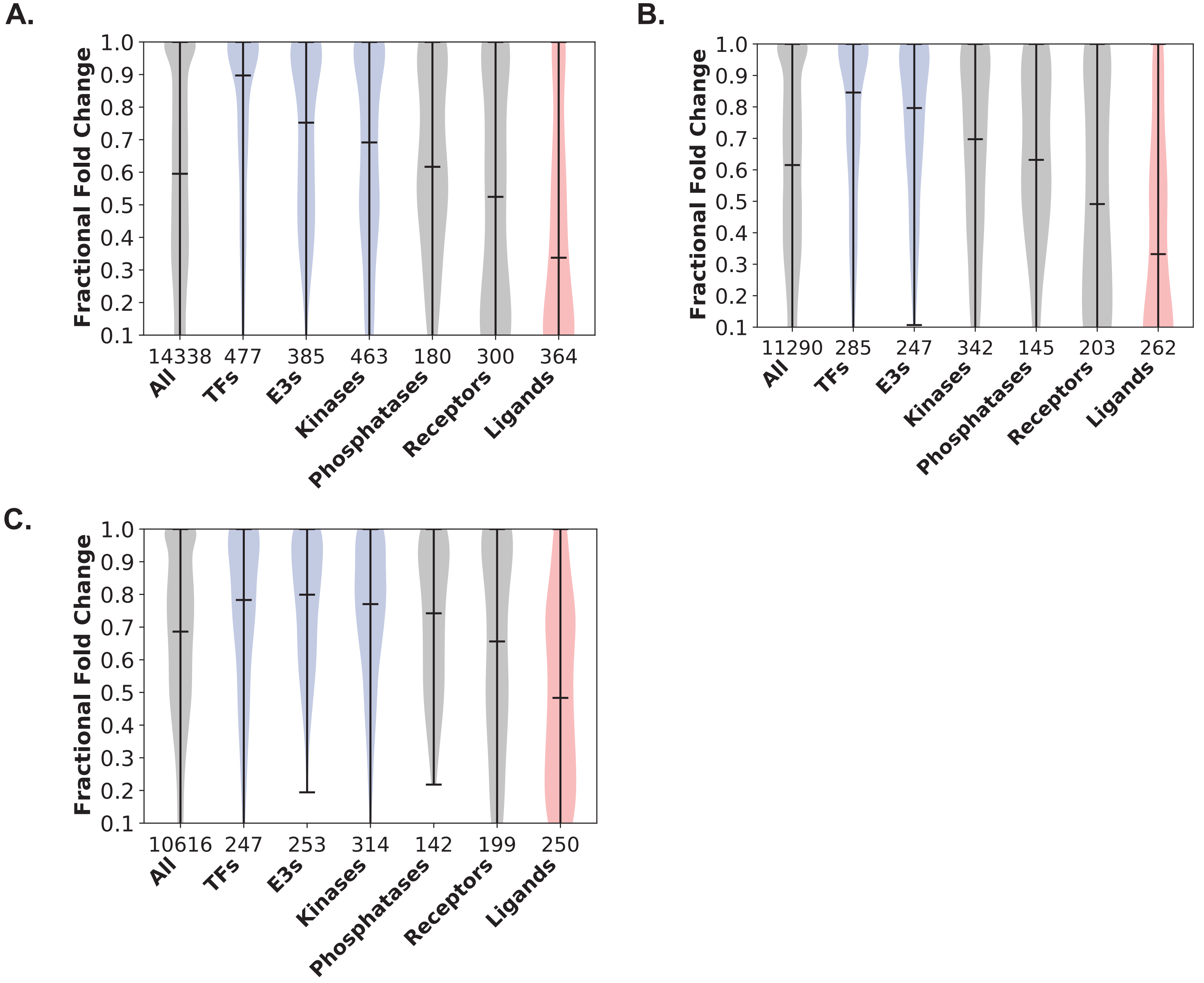
A.
B.
C.
Fractional Fold Change
1.0
0.9
0.8
0.7
0.6
0.5
0.4
0.3
0.2
0.1
14338 477 385 463 180 300 364
11290 285 247 342 145 203 262
10616 247 253 314 142 199 250
All
TFs
E3s
Kinases
Phosphatases
Receptors
Ligands

### Figure S7

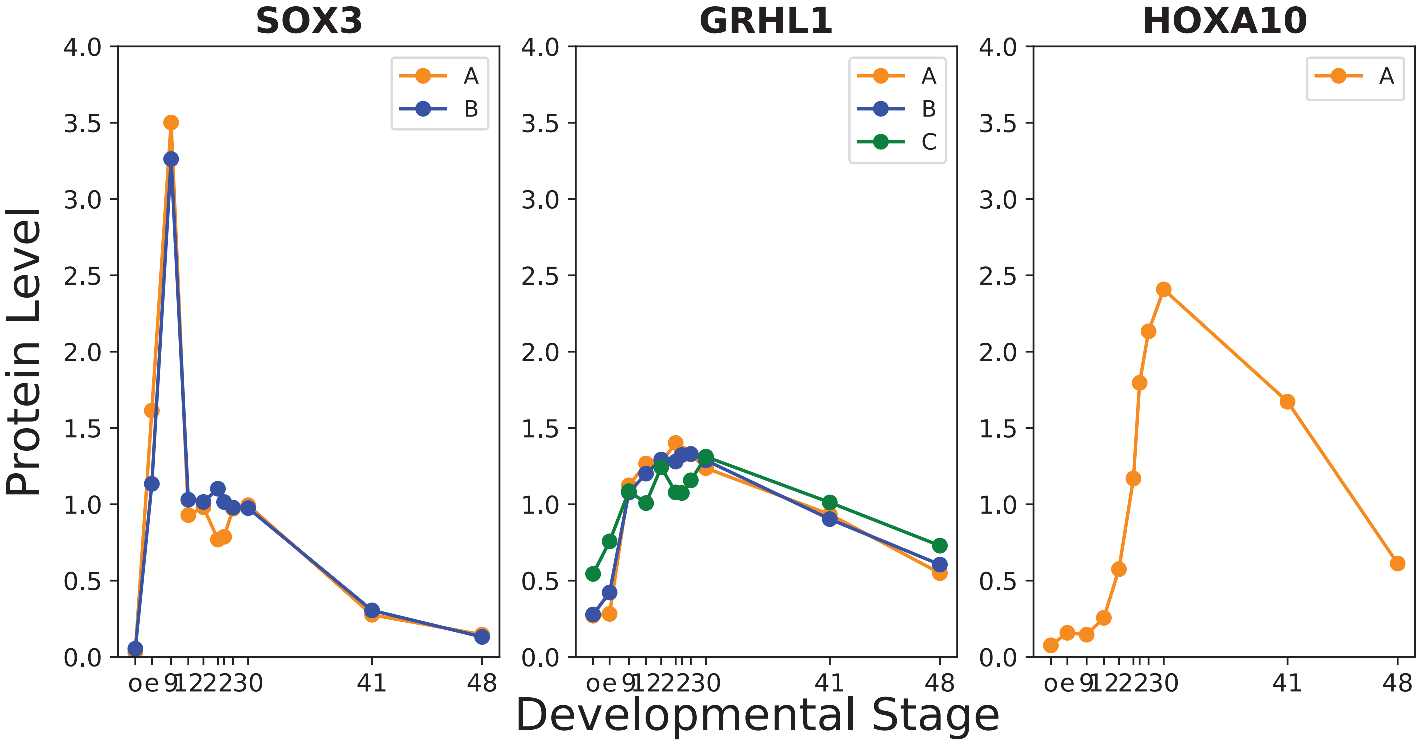

SOX3
GRHL1
HOXA10
Protein Level
Developmental Stage
4.0
3.5
3.0
2.5
2.0
1.5
1.0
0.5
0.0
A
B
C
o e 9 12 22 23 30
41
48

### Figure S8

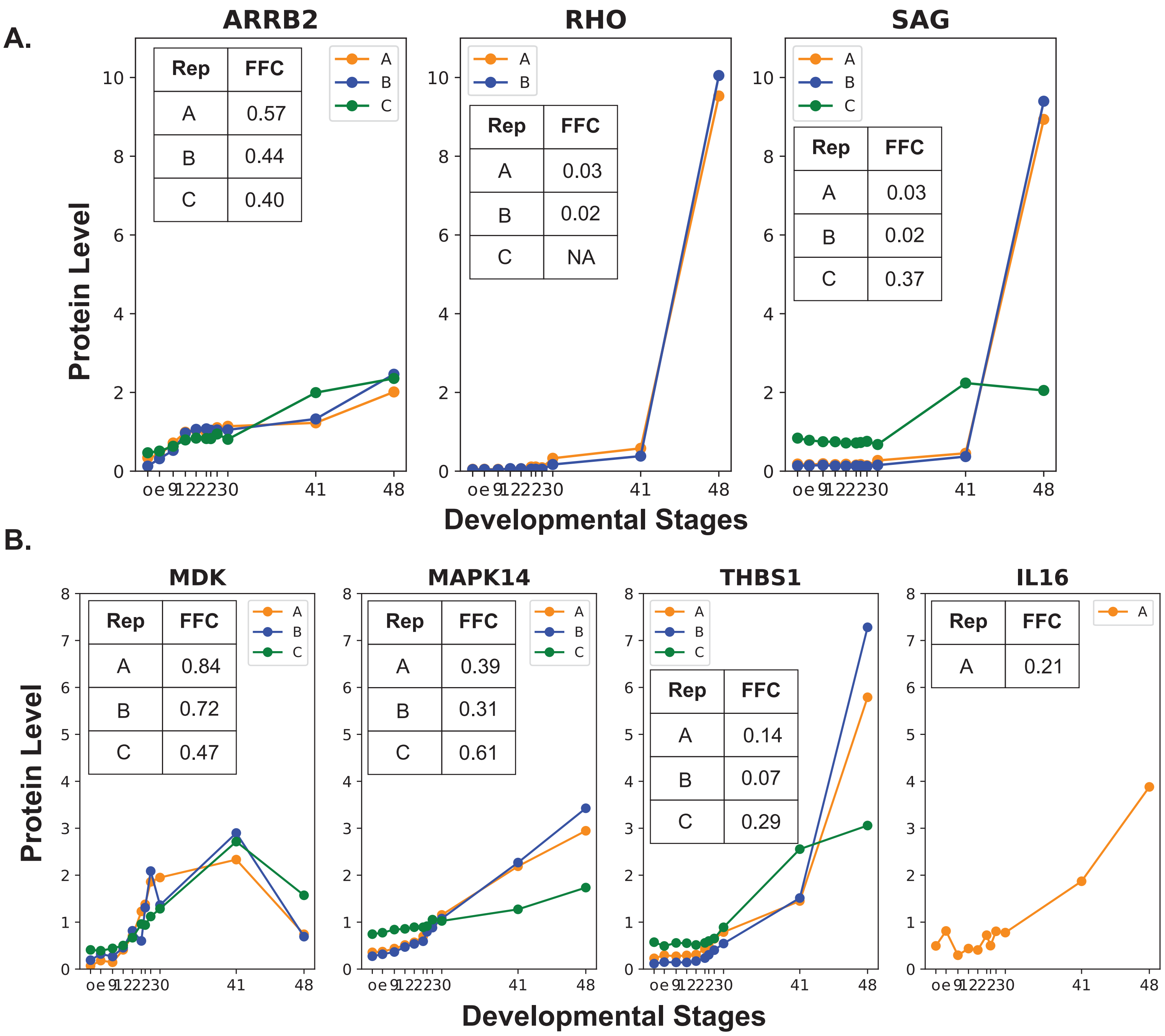
A.
ARRB2
Rep FFC
A 0.57
B 0.44
C 0.40
RHO
Rep FFC
A 0.03
B 0.02
C NA
SAG
Rep FFC
A 0.03
B 0.02
C 0.37
Protein Level
Developmental Stages
o e 9 12 22 30 41 48
B.
MDK
Rep FFC
A 0.84
B 0.72
C 0.47
MAPK14
Rep FFC
A 0.39
B 0.31
C 0.61
THBS1
Rep FFC
A 0.14
B 0.07
C 0.29
IL16
Rep FFC
A 0.21
Protein Level
Developmental Stages

### Figure S9

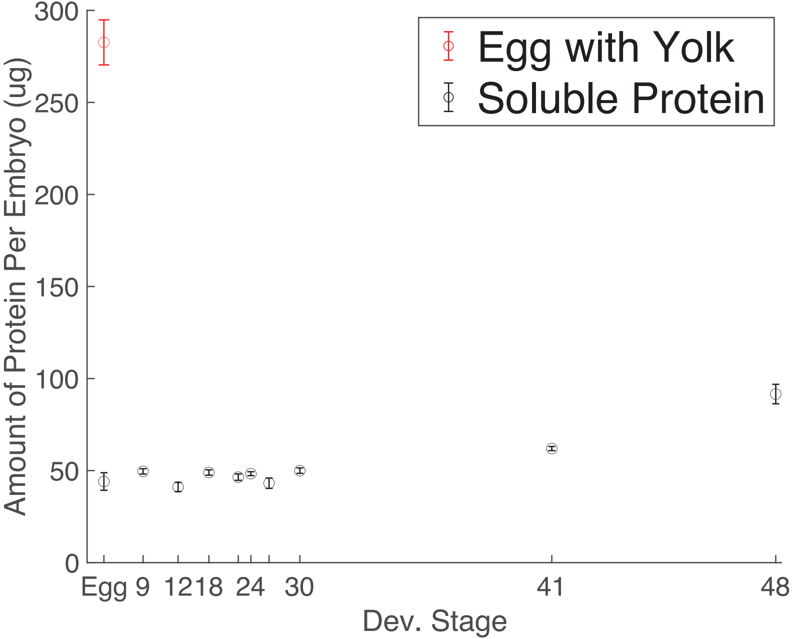
Egg with Yolk
Soluble Protein
Amount of Protein Per Embryo (ug)
300
250
200
150
100
50
0
Egg 9 12 18 24 30 41 48
Dev. Stage

### Figure S10

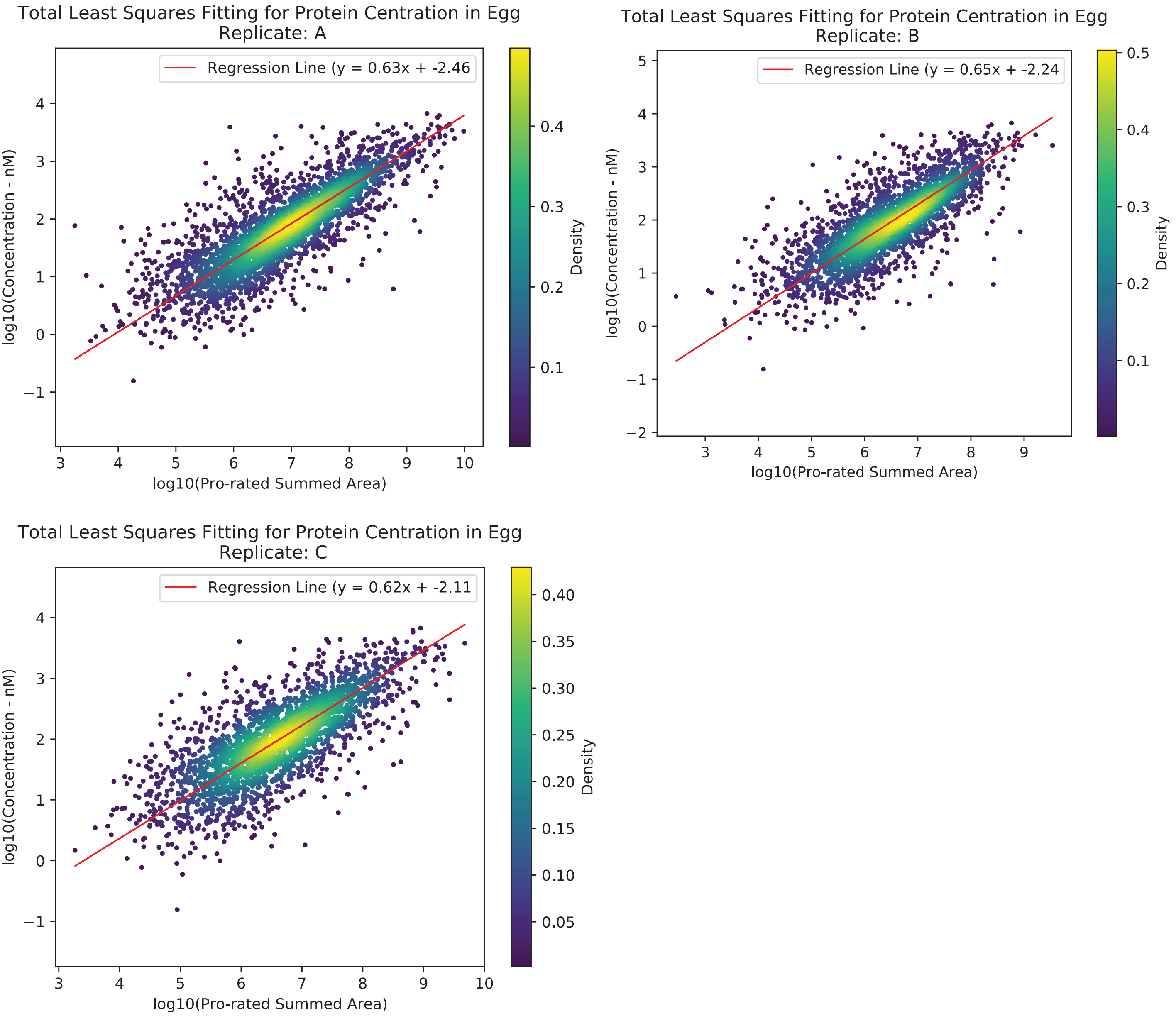
Total Least Squares Fitting for Protein Centration in Egg
Replicate: A
Regression Line (y = 0.63x + -2.46
log10(Concentration - nM)
log10(Pro-rated Summed Area)
Density
Total Least Squares Fitting for Protein Centration in Egg
Replicate: B
Regression Line (y = 0.65x + -2.24
log10(Concentration - nM)
log10(Pro-rated Summed Area)
Density
Total Least Squares Fitting for Protein Centration in Egg
Replicate: C
Regression Line (y = 0.62x + -2.11
log10(Concentration - nM)
log10(Pro-rated Summed Area)
Density

### Figure S11

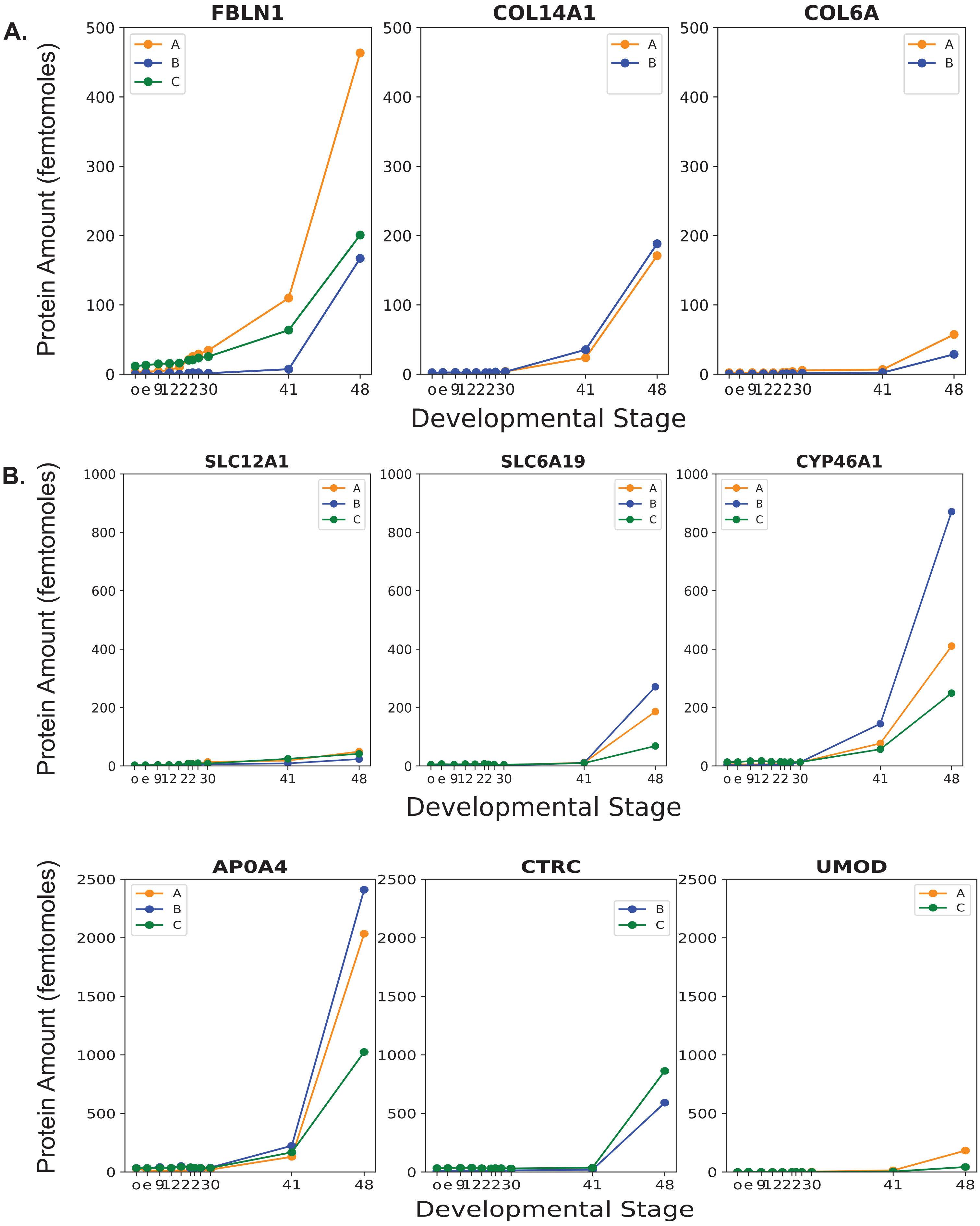

A.
B.
FBLN1
COL14A1
COL6A
SLC12A1
SLC6A19
CYP46A1
AP0A4
CTRC
UMOD
Protein Amount (femtomoles)
Developmental Stage
A
B
C
o e 9 12 22 30
41
48

### Figure S12

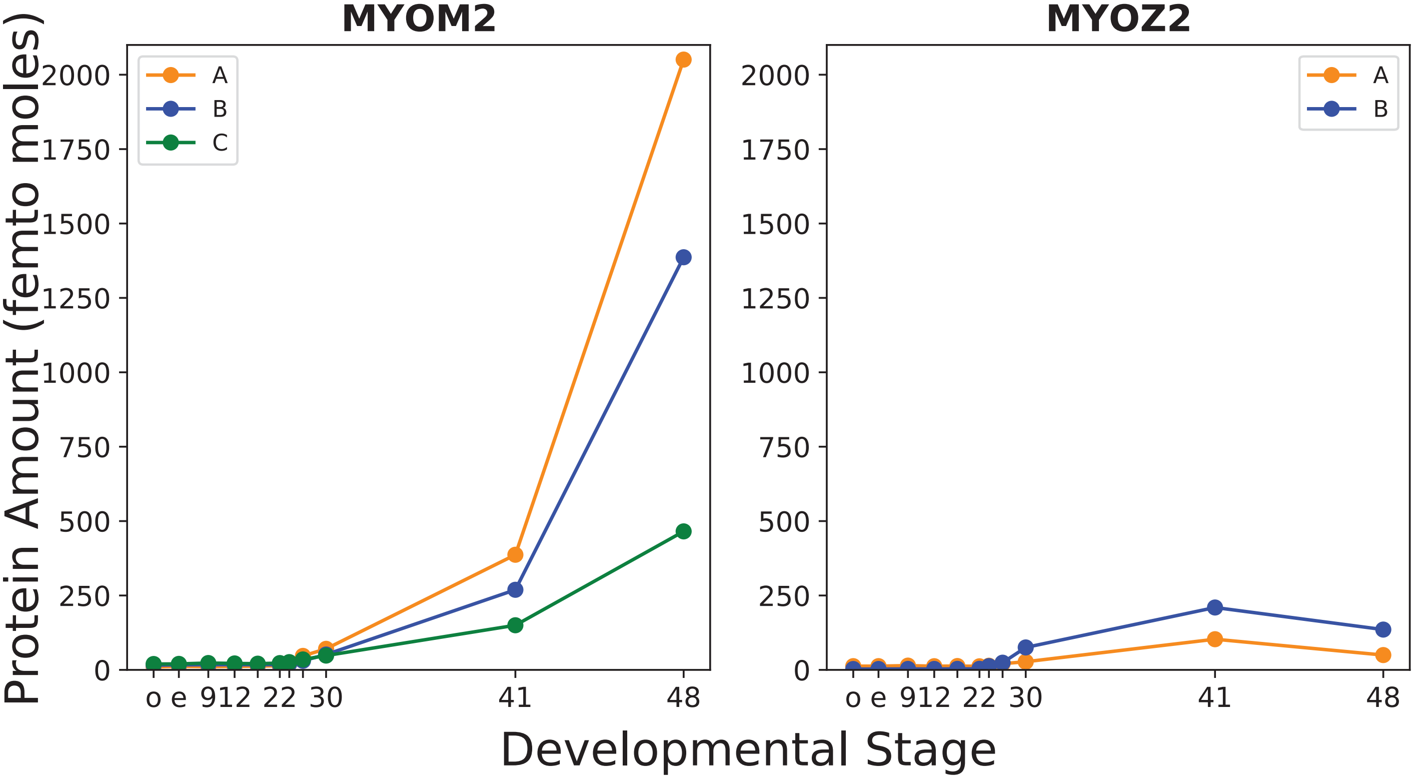
MYOM2
MYOZ2
A
B
C
A
B
Protein Amount (femto moles)
2000
1750
1500
1250
1000
750
500
250
0
o e 912 22 30
41
48
Developmental Stage

#### Figure S13

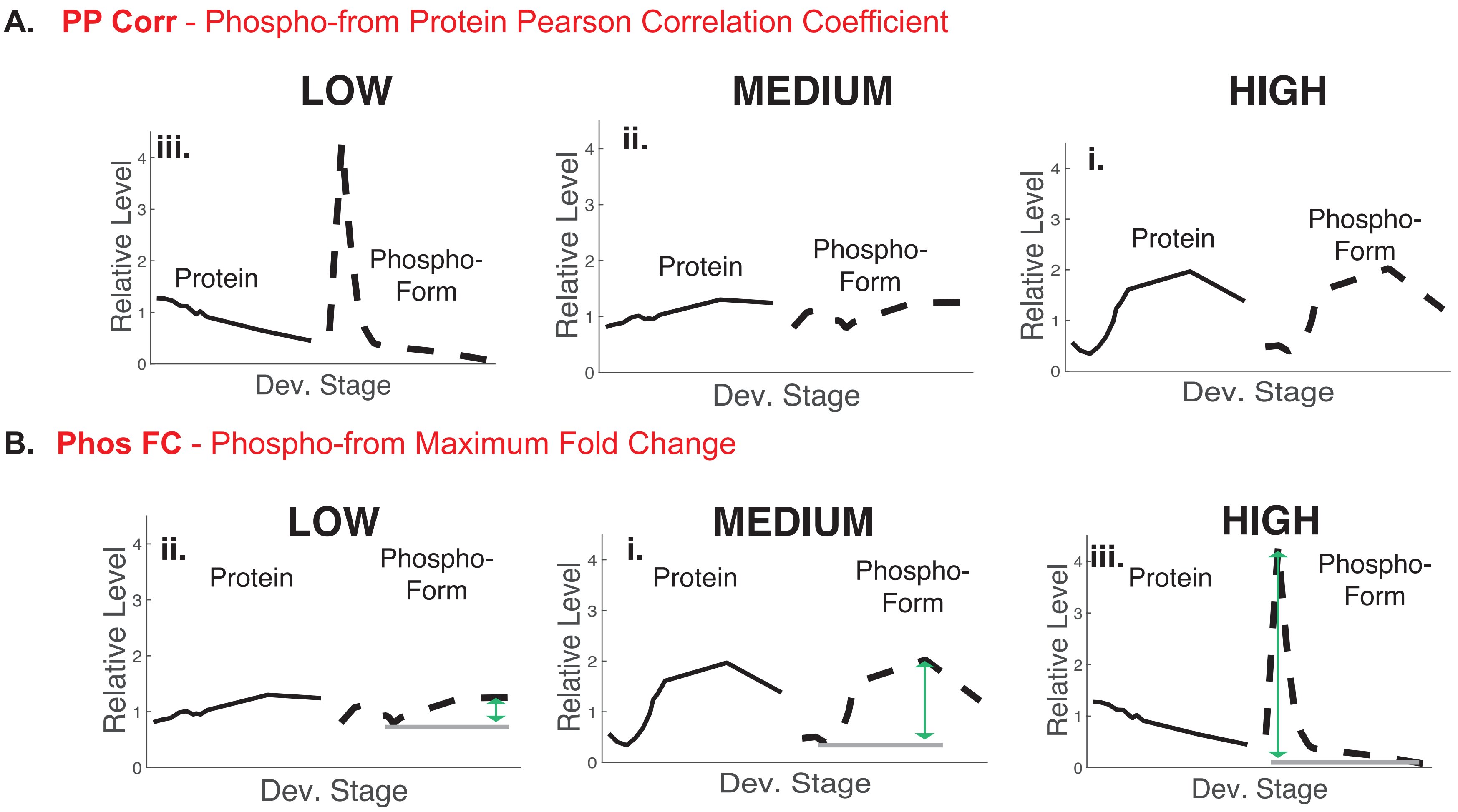

A. PP Corr - Phospho-from Protein Pearson Correlation Coefficient
LOW
iii.
Protein
Phospho-
Form
Relative Level
Dev. Stage
MEDIUM
ii.
Protein
Phospho-
Form
Relative Level
Dev. Stage
HIGH
i.
Protein
Phospho-
Form
Relative Level
Dev. Stage
B. Phos FC - Phospho-from Maximum Fold Change
LOW
ii.
Protein
Phospho-
Form
Relative Level
Dev. Stage
MEDIUM
i.
Protein
Phospho-
Form
Relative Level
Dev. Stage
HIGH
iii.
Protein
Phospho-
Form
Relative Level
Dev. Stage

### Figure S14

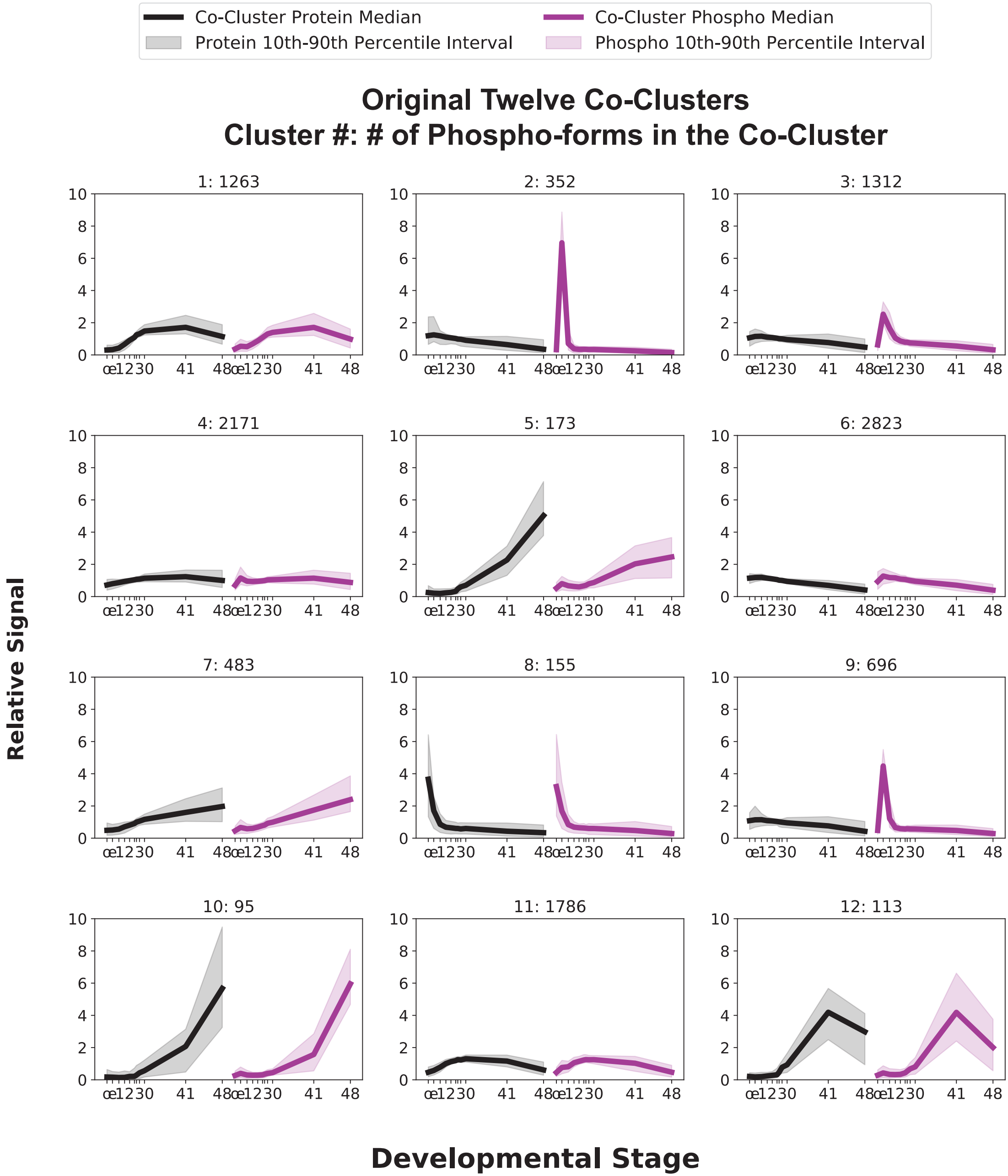
Co-Cluster Protein Median
Protein 10th-90th Percentile Interval
Co-Cluster Phospho Median
Phospho 10th-90th Percentile Interval
Original Twelve Co-Clusters
Cluster #: # of Phospho-forms in the Co-Cluster
1: 1263
2: 352
3: 1312
4: 2171
5: 173
6: 2823
7: 483
8: 155
9: 696
10: 95
11: 1786
12: 113
Relative Signal
Developmental Stage

### Figure S15

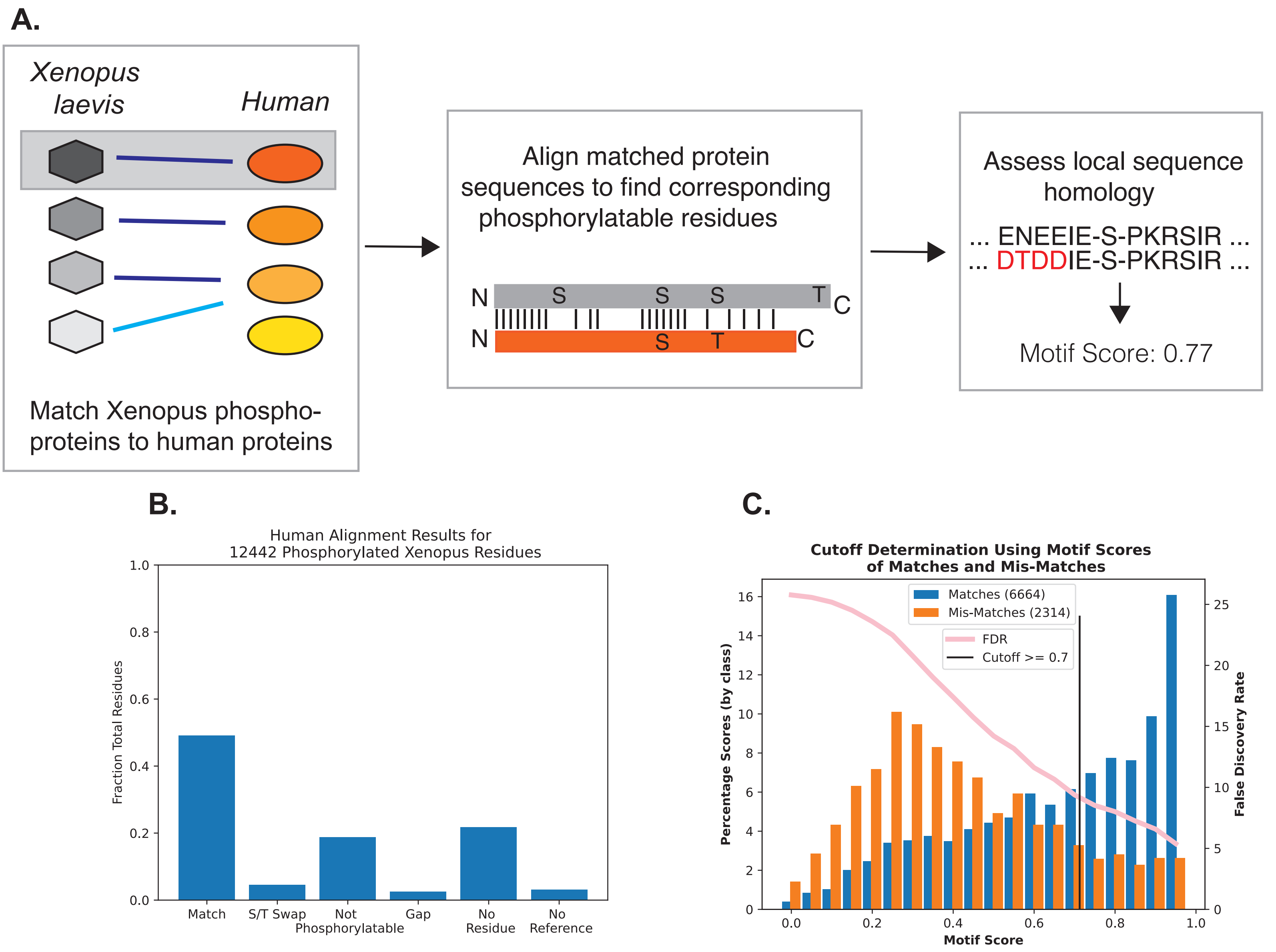
A.
Xenopus laevis
Human
Match Xenopus phospho-proteins to human proteins
Align matched protein sequences to find corresponding phosphorylatable residues
N
S
S
S
T
C
N
S
T
C
Assess local sequence homology
... ENEEIE-S-PKRSIR ...
... DTDDIE-S-PKRSIR ...
Motif Score: 0.77
B.
Human Alignment Results for 12442 Phosphorylated Xenopus Residues
Fraction Total Residues
1.0
0.8
0.6
0.4
0.2
0.0
Match
S/T Swap
Not Phosphorylatable
Gap
No Residue
No Reference
C.
Cutoff Determination Using Motif Scores of Matches and Mis-Matches
Matches (6664)
Mis-Matches (2314)
FDR
Cutoff >= 0.7
Percentage Scores (by class)
16
14
12
10
8
6
4
2
0
False Discovery Rate
25
20
15
10
5
0
0.0
0.2
0.4
0.6
0.8
1.0
Motif Score

### Figure S16

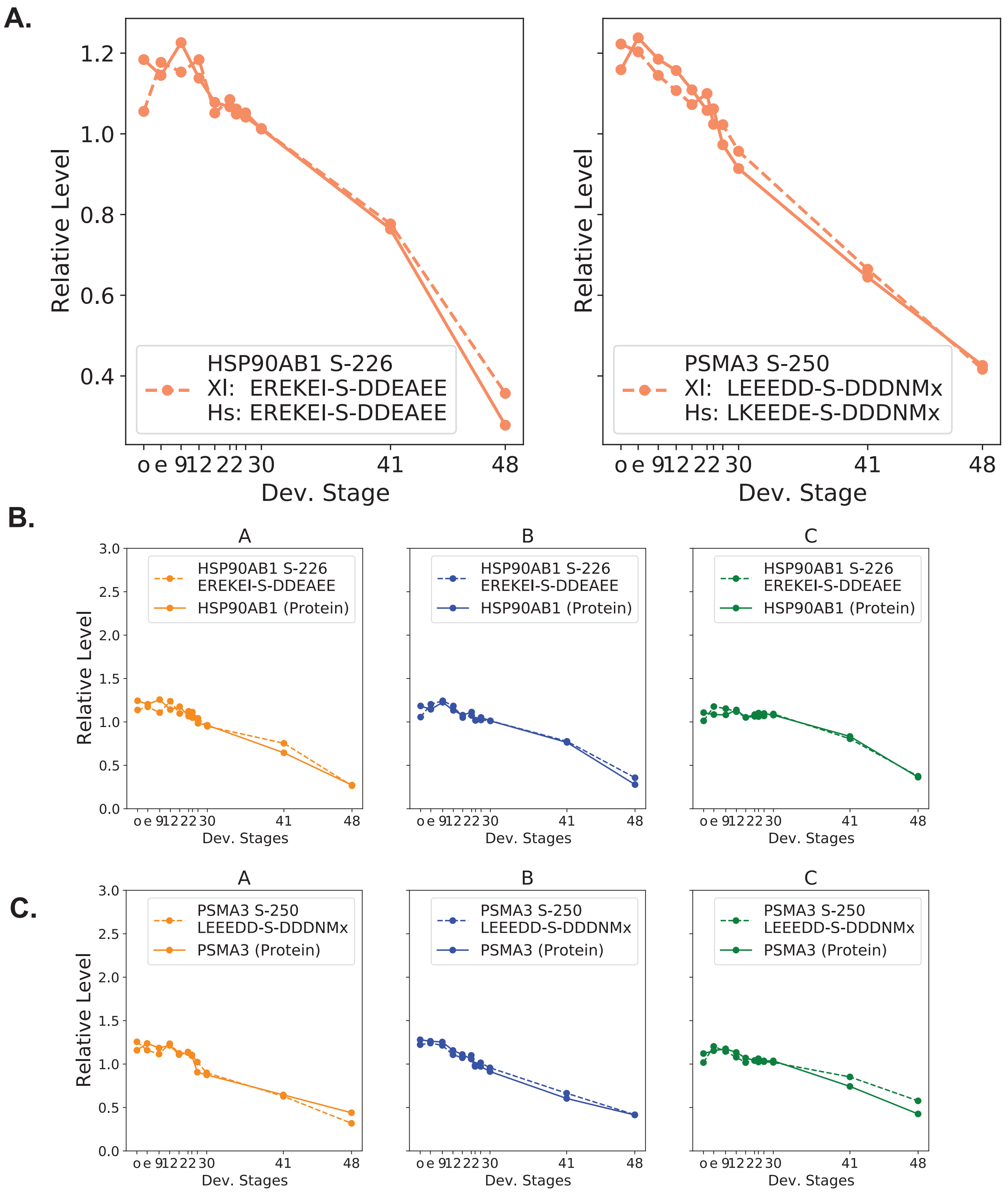
A.
Relative Level
Dev. Stage
HSP90AB1 S-226
Xl: EREKEI-S-DDEAEE
Hs: EREKEI-S-DDEAEE
PSMA3 S-250
Xl: LEEEDD-S-DDDNMx
Hs: LKEEDE-S-DDDNMx
o e 9 12 22 30 41 48
B.
A
B
C
HSP90AB1 S-226
EREKEI-S-DDEAEE
HSP90AB1 (Protein)
Relative Level
Dev. Stages
C.
PSMA3 S-250
LEEEDD-S-DDDNMx
PSMA3 (Protein)

### Figure S17

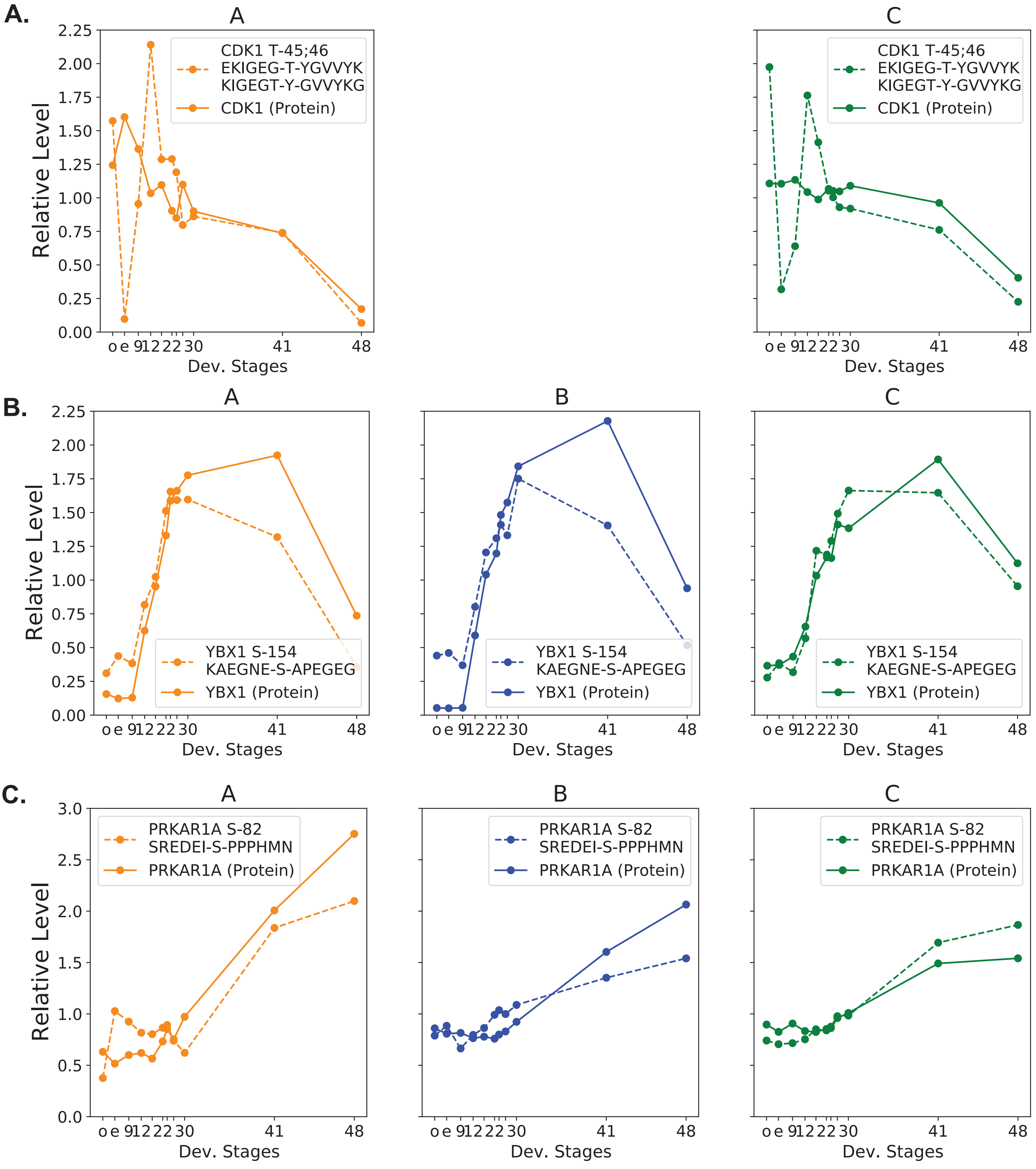
A.
A
C
CDK1 T-45;46
EKIGEG-T-YGVVYK
KIGEGT-Y-GVVYKG
CDK1 (Protein)
Relative Level
Dev. Stages
o e 9 12 22 30 41 48
B.
A
B
C
YBX1 S-154
KAEGNE-S-APEGEG
YBX1 (Protein)
Relative Level
Dev. Stages
o e 9 12 22 30 41 48
C.
A
B
C
PRKAR1A S-82
SREDEI-S-PPPHMN
PRKAR1A (Protein)
Relative Level
Dev. Stages
o e 9 12 22 30 41 48

### Figure S18

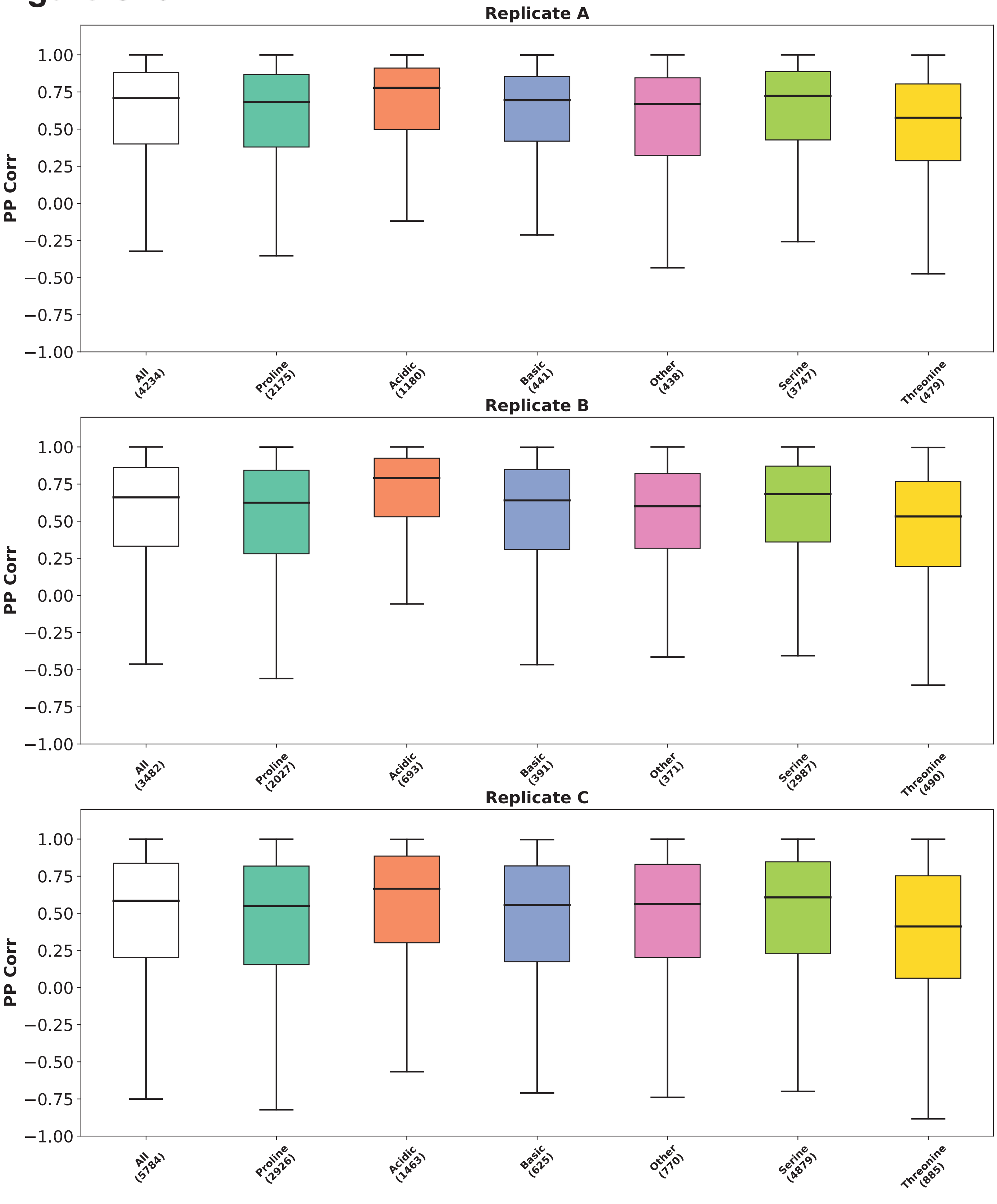
Replicate A
PP Corr
1.00
0.75
0.50
0.25
0.00
−0.25
−0.50
−0.75
−1.00
All (4234)
Proline (2175)
Acidic (1180)
Basic (441)
Other (438)
Serine (3747)
Threonine (479)
Replicate B
All (3482)
Proline (2027)
Acidic (693)
Basic (391)
Other (371)
Serine (2987)
Threonine (490)
Replicate C
All (5784)
Proline (2926)
Acidic (1463)
Basic (625)
Other (770)
Serine (4879)
Threonine (885)

### Figure S19

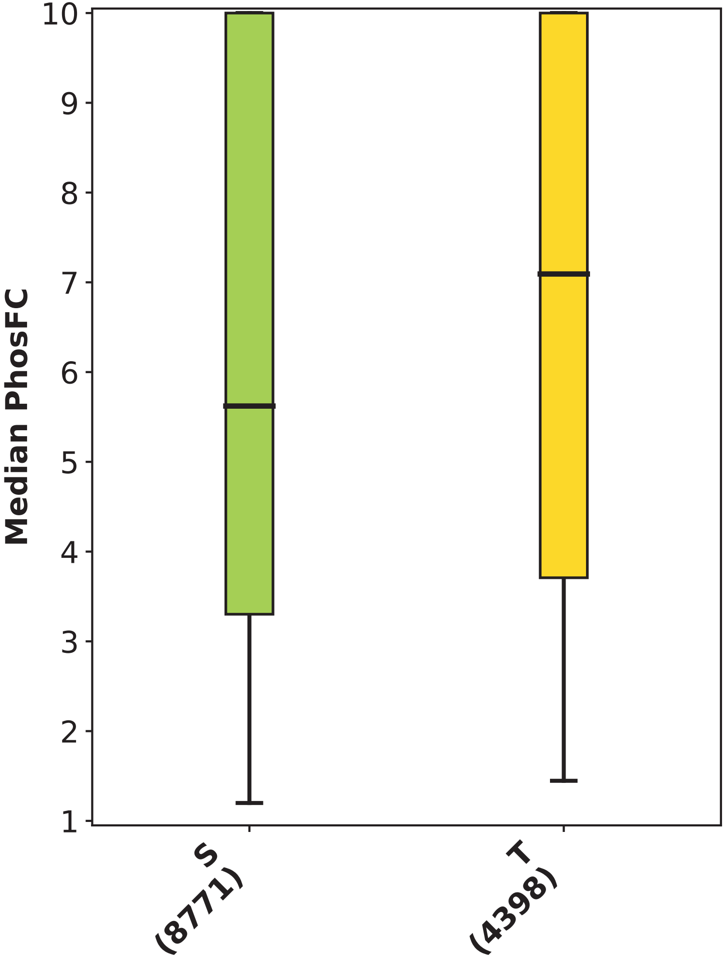

### Figure S20

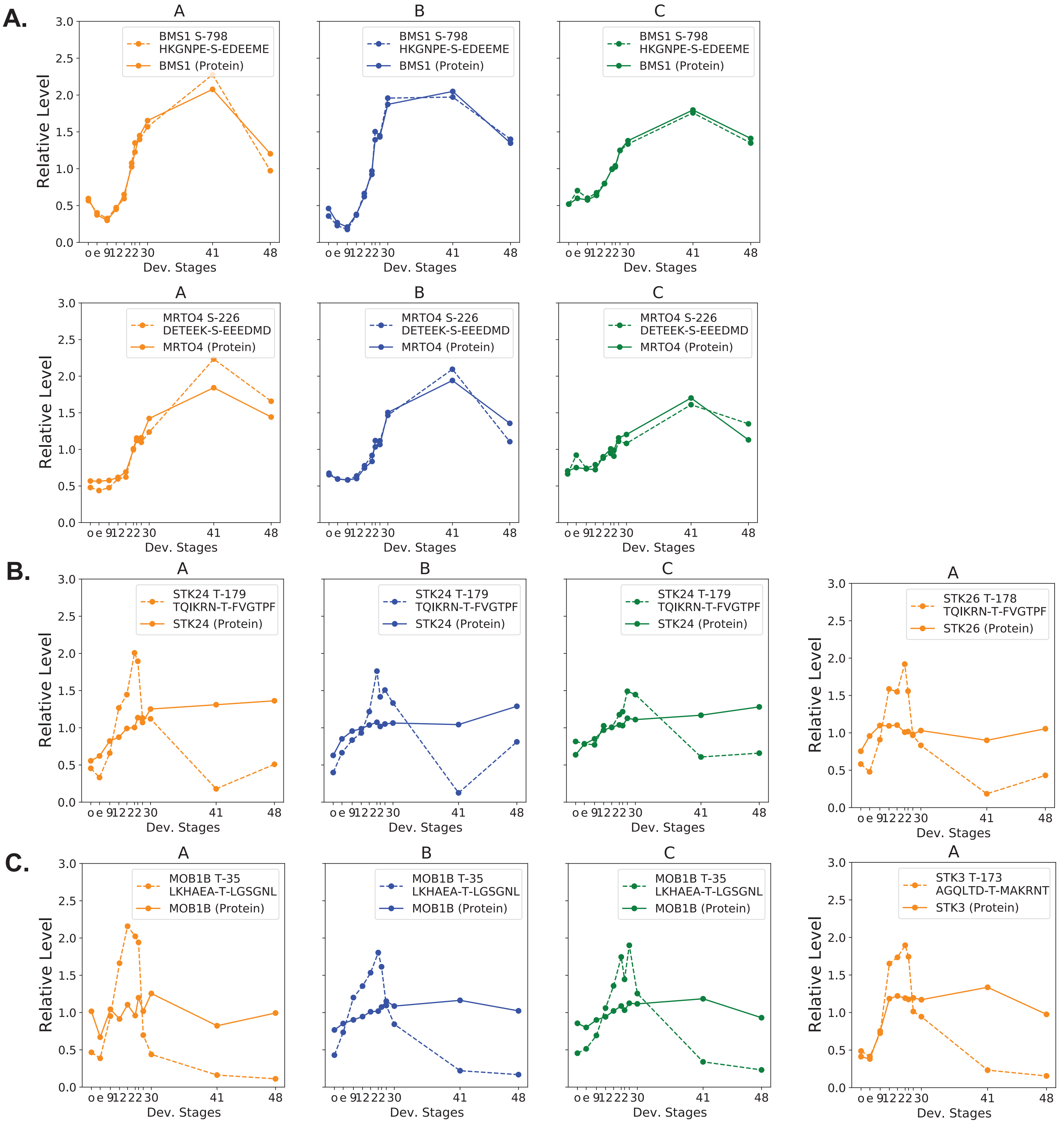

A.
A
BMS1 S-798
HKGNPE-S-EDEEME
BMS1 (Protein)
B
BMS1 S-798
HKGNPE-S-EDEEME
BMS1 (Protein)
C
BMS1 S-798
HKGNPE-S-EDEEME
BMS1 (Protein)
A
MRTO4 S-226
DETEEK-S-EEEDMD
MRTO4 (Protein)
B
MRTO4 S-226
DETEEK-S-EEEDMD
MRTO4 (Protein)
C
MRTO4 S-226
DETEEK-S-EEEDMD
MRTO4 (Protein)
B.
A
STK24 T-179
TQIKRN-T-FVGTPF
STK24 (Protein)
B
STK24 T-179
TQIKRN-T-FVGTPF
STK24 (Protein)
C
STK24 T-179
TQIKRN-T-FVGTPF
STK24 (Protein)
A
STK26 T-178
TQIKRN-T-FVGTPF
STK26 (Protein)
C.
A
MOB1B T-35
LKHAEA-T-LGSGNL
MOB1B (Protein)
B
MOB1B T-35
LKHAEA-T-LGSGNL
MOB1B (Protein)
C
MOB1B T-35
LKHAEA-T-LGSGNL
MOB1B (Protein)
A
STK3 T-173
AGQLTD-T-MAKRNT
STK3 (Protein)
Relative Level
Dev. Stages
o e 9 12 22 30 41 48
0.0 0.5 1.0 1.5 2.0 2.5 3.0
